# 34-parameter full spectrum immunophenotyping panel of human regulatory and effector lymphocytes

**DOI:** 10.1101/2024.03.21.585958

**Authors:** Zoya Georgieva, Valerie Coppard, Jennie HM Yang, Richard Grenfell, Joanne Jones

## Abstract

This 34-marker sentinel, intracellular, full-spectrum flow cytometry panel profiles regulatory and effector T, B and NK lymphocytes in human cryopreserved peripheral blood mononuclear cells. The panel focuses on cell trafficking, activation, exhaustion and proliferation, and permits easy customisation in two positions to accommodate other targets of the user’s interest. By combining breadth and depth of phenotyping, this panel is designed to maximise the information obtained from limited cell material and therefore will be particularly useful in mechanistic studies of immunomodulatory drugs for autoimmune disease, cancer and transplantation, where multiple immune populations may be affected.

## Background

A breakdown in suppressive mechanisms leads to autoimmunity, while excessive immune suppression or effector cell exhaustion are hallmarks of cancer. Immunotherapies which target these defects may act on both regulatory and effector lymphocyte populations due to the highly overlapping expression of their molecular targets.

Interleukin-2 (IL-2) is an exemplar of this principle. At high doses, IL-2 enhances T-effector (Teff) anti-tumour immunity, but its effectiveness is limited by the concomitant expansion of the highly IL-2 responsive CD4^+^CD127^lo^CD25^hi^FOXP3^+^ T-regulatory cells (Tregs)(1). It is now known that IL-2 is non-dispensable for Treg function, and at low doses, it specifically promotes immune tolerance by expanding Tregs, as well as CD56^bright^ NK cells and CD25+ IL-10 producing B-regulatory cells (Bregs)(2), while diminishing Th1, Th17 and IL-21-producing Tfh effectors(3)(4). Another widely known example is that immune checkpoint blockade (ICB), which aims to enhance anti-tumour cytotoxic function, also may induce iatrogenic autoimmunity through its action on Tregs.

Given these potential pleiotropic effects, when developing novel immunotherapies, it is important to examine the impact of the drug on both regulatory and effector cells in multiple lymphocyte subpopulations.

CD4^+^ Tregs are the most extensively studied regulatory population and are known to be dysfunctional in human autoimmunity, such as systemic lupus erythematosus(5), type 1 diabetes(6), and multiple sclerosis(7). They also play a role in tumour immune evasion(8). Their lesser-known CD8 counterparts (FOXP3^+^ CD8^+^ Tregs) have also been shown to be enriched in tumour sites(9), and we recently demonstrated that they can be isolated from healthy human non-lymphoid tissue (10), raising questions about their physiological role and therapeutic potential. B cells with regulatory function have also been described including: CD24^hi^CD38^hi^ B cells which are reported to be defective in SLE(11)); CD24^hi^CD27^+^B cells, which are reduced in chronic graft-versus-host disease (12) and expanded in bullous pemphigoid(13); and adenosine-producing CD39^+^ B-cells, which play a role in modulating rheumatoid arthritis activity(14)). Finally CD56^bright^ NK cells, which are often thought of as regulatory, expand in response to some immunomodulatory treatments for multiple sclerosis(15). CD56^bright^ NK cells can preferentially kill proliferating autoreactive CD4 Teffs that have escaped negative selection(16).

Since T, B and NK lineages all contain lymphocytes with regulatory and effector properties, the ability to achieve comprehensive phenotyping with limited cellular material would be a great advantage in determining the impact of a novel immunotherapy.

To achieve this, we designed a 34-marker spectral cytometry panel, which juxtaposes selected markers involved in the function of regulatory and effector T, B and NK lymphocytes. We prioritised the inclusion of highly co-expressed surface and intracellular markers with well-characterised roles in the T-cell compartment, the main focus of the panel. Many of these markers are also expressed in B- and NK-cells, where their relevance to disease is emerging. Therefore, the use of this panel could also lead to novel insights into altered immune phenotypes during immunomodulatory therapy.

The panel characterizes naïve and memory CD4 and CD8 T-cells, CD4 and CD8 FOXP3+ T-regulatory cells, three NK cell subsets, two primary NKT cell subsets, and twelve B-cell subsets (Figure 1 and Supplementary Figure 1). Viability stain and CD14 exclude dead cells and monocytes. Two positions on the panel can be customised with user-selected targets.

**Figure 1.**
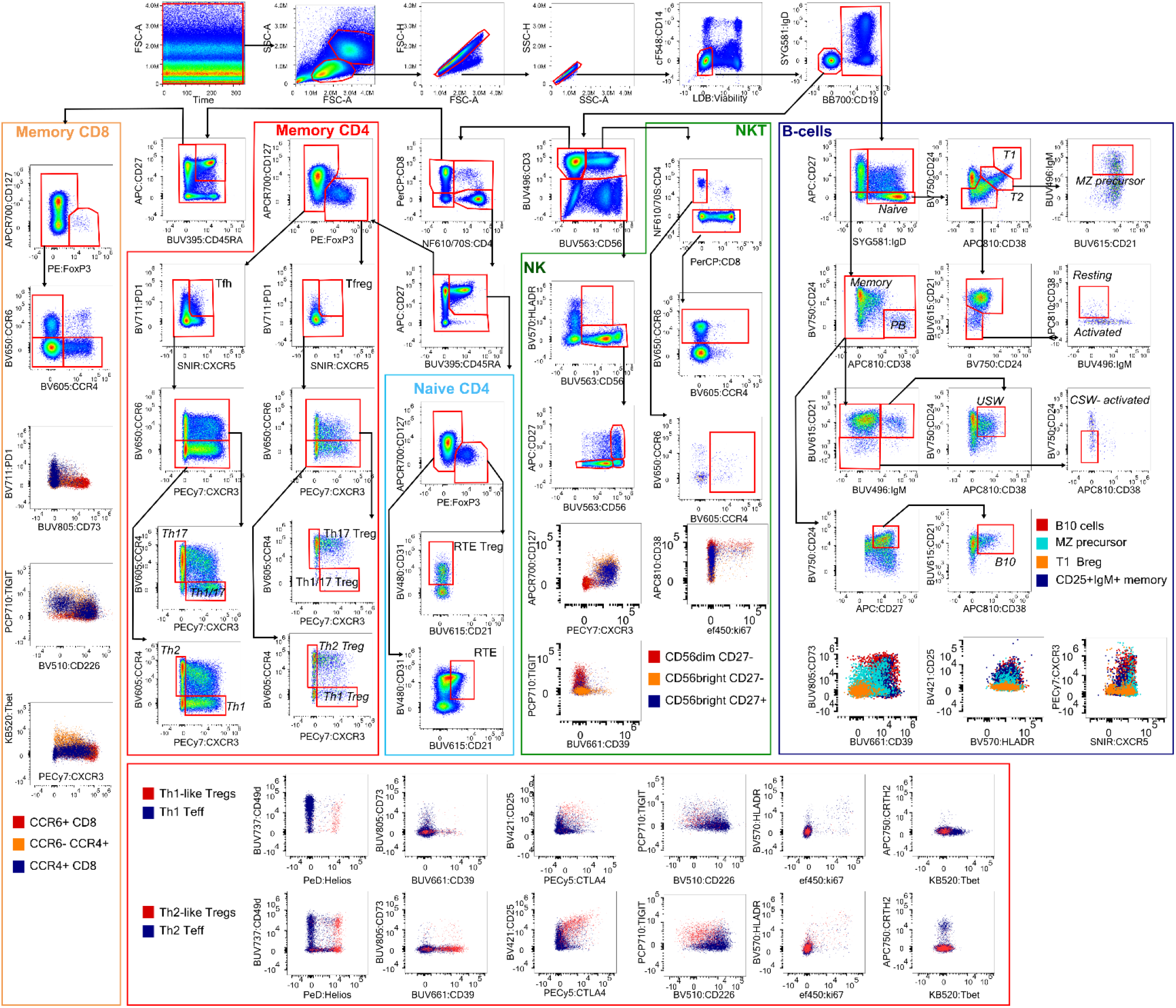
Gating strategy of one healthy control sample. Monocytes and dead cells are excluded first, followed by onvolution of B- and T-cells (IgM and CD3 share a fluorochrome). The gating hierarchy shown is one of several sible strategies. Colour dot plots show the expression of selected tertiary markers on the lymphocyte population cated in nearby colour legends (e.g. CCR6+ CD8 T-cells; T1 B-cells, CD56^bright^ NK cells, Th1-like Tregs and T-effectors etc).

CD3 and IgM share the BUV496 fluorochrome in a sentinel approach. Initial gating separates CD19^+^ from CD3^+^ cells, followed by CD56^+^NK and CD56^+^CD3^+^ NKT cells. CD45RA and CD27 discriminate between naïve and memory T-cells. Regulatory T-cells are identified by CD127 and FOXP3. The use of 2 clones for FOXP3 is recommended to capture total Tregs more efficiently(17) and we applied this principle of using two clones binding different epitopes to CD25 staining (18). The transcription factor HELIOS was also included as a marker of Treg phenotypic stability under inflammatory conditions(19). Our panel was also designed to be equally effective at identifying Tregs using two other commonly employed gating strategies – i.e. FOXP3/HELIOS and CD127^lo^/CD25^hi^.

Memory CD4 and CD8 T-effector cells become polarised into two major subsets (among many others) producing predominantly type 1 (IL-2, IFN-γ, TNF-α) or type 2 (IL-4, IL-5) cytokines, with corresponding chemokine receptor profiles, which govern the immune surveillance of tissues(20). Tregs emulate the expression of the same chemokine receptors to navigate to inflammatory sites(21). Our panel is able to identify the following subsets of polarised Teffs and Tregs: CXCR5^+^PD1^hi^Tfh cells; CCR6^−^ CCR4^−^CXCR3^+^ Th1; CCR6^−^CCR4^+^CXCR3^−^ Th2; CCR6^+^CCR4^+^CXCR3^−^ Th17; And CCR6^+^CCR4^−^ CXCR3^+^ Th1/17 in CD4 memory T-cells. Intracellular TBET (which governs CXCR3 and IFNγ expression) is optionally included for confirmation of Th1 commitment(22), while the surface receptor CRTH2 indicates Th2 commitment(23).

Additionally, the panel identifies skin-tropic CCR4^+^ type 1/type 2 cytokine producers (24) and mucosal-homing CCR6^+^ (25)) CD8 T-effectors. TIGIT/CD226, CD49d, CD39/CD73, CD70, CTLA-4 and CD21/CD31 have been included due to their well-characterised roles in T-cells, particularly in autoimmune disease and cancer, summarised in Supplementary Table 1.

Human NK cells can be classified as (i) CD27^−^CD11b^−^ (which are mostly CD56^bright^, and are sometimes called “tolerant” due to their low degranulation propensity), (ii) CD27^+^CD11b^+/−^ (which are also CD56^bright^ and may be considered “regulatory”, as they eliminate pro-inflammatory T-cells) and (iii) CD27^−^CD11b^+^ (mostly overlapping the large CD56^dim^CD16^−^ subset which is predominantly “cytotoxic”)(26). Our panel delineates three NK cell populations without the use of CD11b: CD56^bright^CD27^−^, CD56^bright^CD27^hi^ NK cells, and CD56^dim^CD27^−^. Published evidence for the roles of CXCR3, CD226, TIGIT, PD-1, CD39 and TBET in NK cells is summarised in Supplementary Table 2.

Key stages of B-cell maturation can be identified according to a gating convention using CD27, IgD, IgM, CD24, CD38 and CD21(27). Bregs do not have lineage-defining markers, and are enriched in multiple B-cell populations, including transitional CD24^hi^CD38^hi^(11), memory IgM^+^ (28), and CD24^hi^CD27^hi^ (“B10”)(12). Although Bregs are best identified based on IL-10 production, this panel is not designed for use on stimulated cells, since this would alter the *ex vivo* expression patterns of numerous other markers. The function of CD39, CD73, chemokine receptors, TIGIT and TBET are of potential interest in the human B-cell compartment, as summarised in Supplementary Table 3.

FMO controls are recommended for the following markers on this panel: CD21 (in T-cells), CD226, PD-1, CXCR5, CD24, CD70, CD38, CTLA-4 and TBET.

Two markers on this panel (TBET Kiravia Blue 520, and CRTH2 APCFire750) can be substituted with user-selected alternatives. The B1/2 and R7 channels were selected due to the wide availability of bright fluorochromes reagents with narrow primary emission peaks, and the relatively limited impact of other markers in this region. In principle, this permits straightforward direct substitution of most markers regardless of their expression pattern. For example, in order to validate expression data pertaining to the Treg compartment in alemtuzumab-treated individuals with MS, we substituted GITR (BB515) and OX-40 (APC-Fire750). Sample data are shown in Supplementary Figure 2. This substitution did not require any additional pilot experiments, which was advantageous due to our lack of redundant patient samples. Nevertheless, a validation experiment would still be recommended if the researcher has no prior experience of the marker or study population.

In summary, we have developed a 34-marker sentinel full-spectrum panel, which characterises major populations of regulatory lymphocytes and their effector cell counterparts in human peripheral blood. Our panel addresses a need for high-resolution in-depth phenotyping of co-expressed molecular targets and is, to our knowledge, the first panel specifically designed for full-spectrum cytometry that incorporates intracellular staining of human cells. Going considerably beyond enumeration of population frequencies, this panel conveys detailed phenotypic information and provides an opportunity for novel biomarker discovery.

### Similarity to other OMIPs

Some markers are shared with OMIPs: -04, -06, -15, -17, -18, -53, which characterise Tregs and/or T-effectors; -47 which characterises B-cells, and -80 which examines immune reconstitution after stem cell transplant. This panel has two key unique features: a) high resolution of co-expressed tertiary markers on 3 different lymphocyte lineages; b) inclusion of intracellular targets, which are not yet featured in other published spectral panels.

**Table 1.**
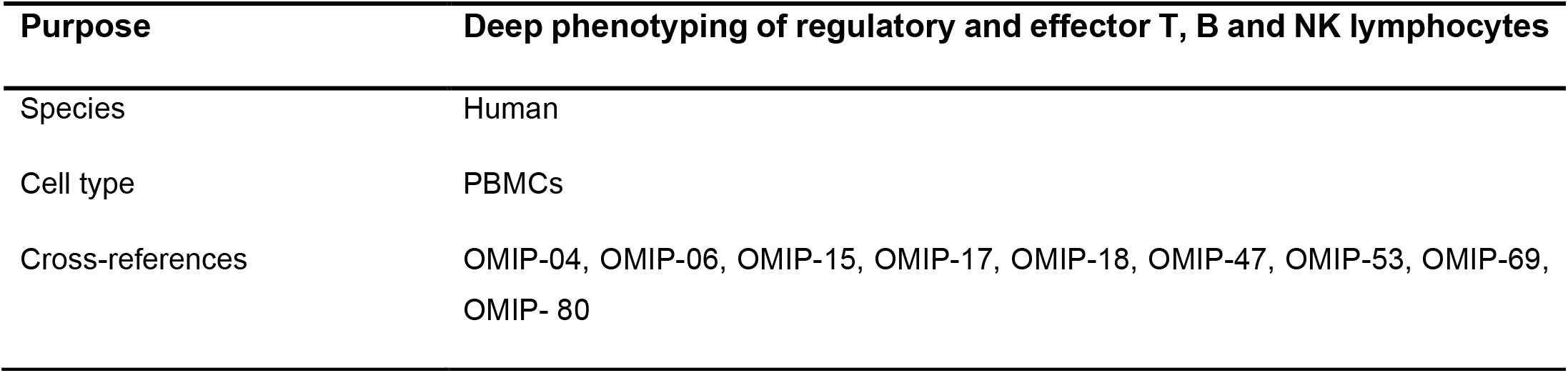
Summary characteristics of OMIP-xx.

**Table 2.**
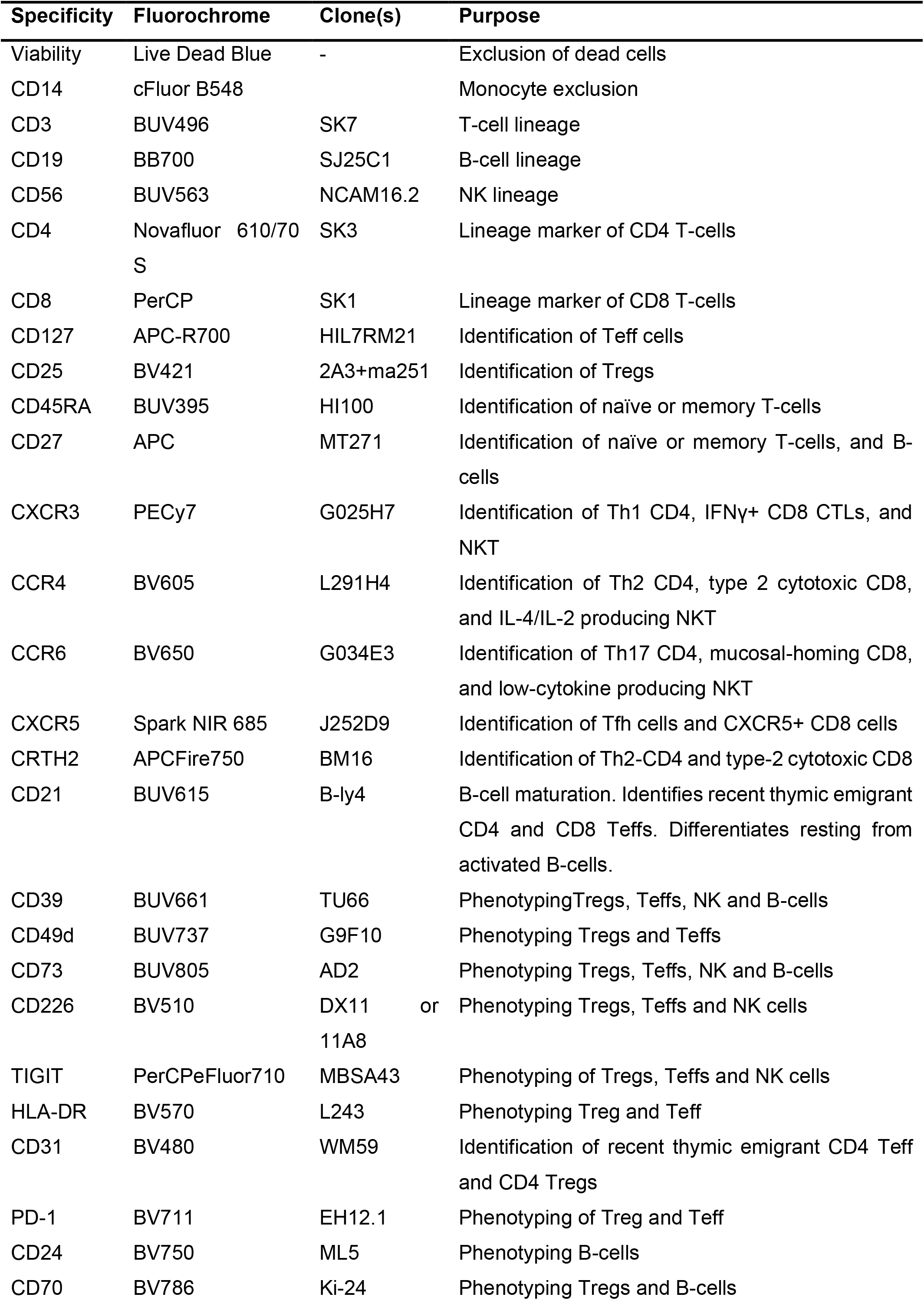
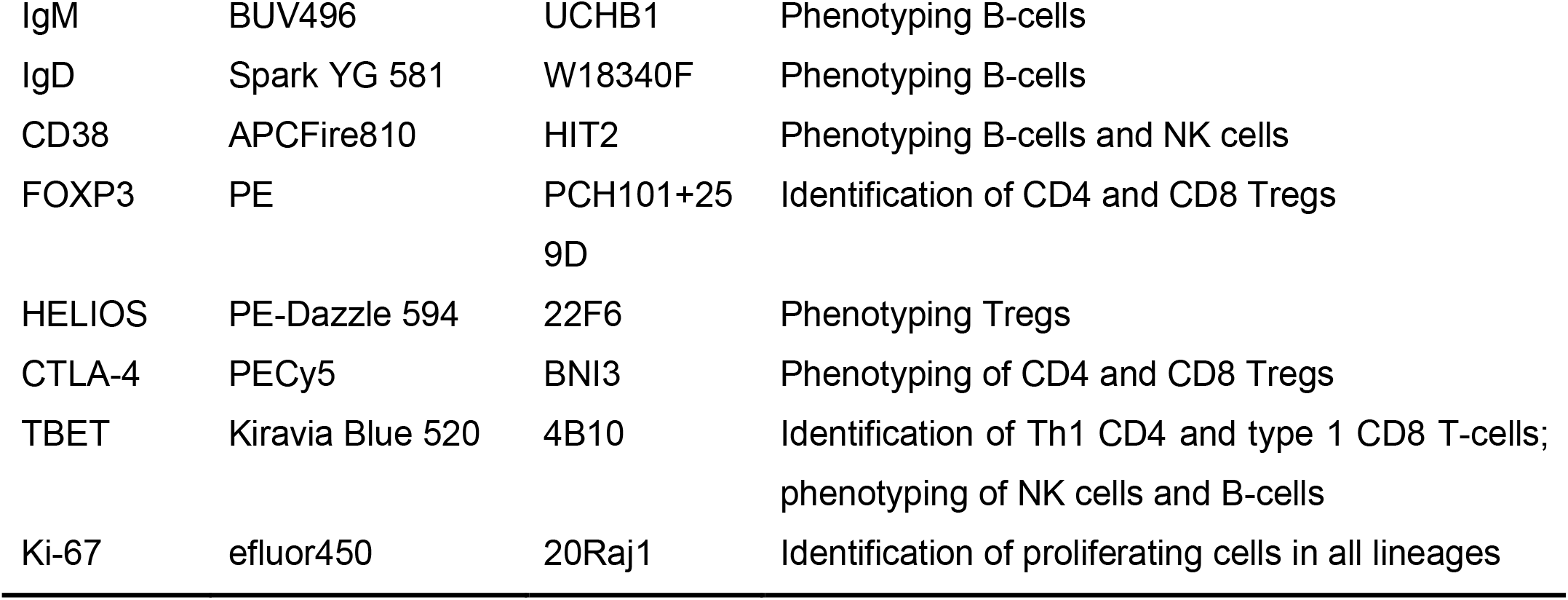
Reagents used for OMIP-xx.

### Ethics

Human PBMCs were prepared from healthy control volunteers and patients with relapsing remitting multiple sclerosis, who had given free informed consent under ethics REC-11/33/0007.

## Supporting information

Supplemental information: panel development, materials, protocols & example use

## Author contributions

Zoya Georgieva-conceptualization; data curation; formal analysis; methodology; validation; visualization, writing ‐original draft; writing ‐review and editing

Valerie Coppard-conceptualization; data curation, writing ‐review and editing Jennie Yang-conceptualization, methodology

Richard Grenfell-methodology, resources, supervision, software, writing-review and editing Joanne Jones-project administration; resources; supervision; writing ‐review and editing

