## Supplemental information: panel development, materials, protocols & example use for "34-parameter full spectrum immunophenotyping panel of human regulatory and effector lymphocytes"

### 1. Supplementary information- Panel development

#### 1.1. Initial fluorochrome selection

The similarity index (SI) and complexity index (CI) were used to guide reagent choice for the panel. Commonly available fluorochromes with unique spectra were selected first, then candidate fluorochromes with SI >0.8 were considered for compatibility with these and selected to minimise impact on the CI. Highly similar fluorochromes were assigned to minimise the probability of co-expression, according to the Astrolabe mass cytometry data set(29). We did not use custom-made reagents or self-conjugated fluorochromes to achieve maximum accessibility and reproducibility. The published normalised spectra and similarity indices for the final selected fluorochromes are shown in Supplementary Figure 3 and Supplementary Figure 4.

#### 1.2. Allocation of primary lineage-identifying markers

Lineage-identifying markers were allocated to highly similar fluorochromes without loss of resolution, since they have predictable binary patterns of expression, as follows:

- CD3 to BUV496, which spreads very little into any other channels but receives substantial spread from UV-range autofluorescence in fixed permeabilised cells. Therefore, this fluorochrome was only suitable for a densely expressed lineage marker such as CD3.

- CD56 to BUV563, which is not co-expressed with PE (FOXP3), optimising resolution of Tregs.

- CD4 to Novafluor610/70S. This fluorochrome has a unique spectrum with smaller secondary emission peaks than other blue-laser excitable dyes. Although it is said to aggregate easily, we experienced no problems after centrifuging the new vial at 14,000G (4C$^{\circ}$) before first use.

- CD8 to PerCP, a very dim fluorochrome which minimises spread from its secondary emission peaks.

- CD19 to BB700. This fluorochrome has high quantum yield, fully separating $CD19^{+}$ from $CD19^{-}$ cells, which was essential in our sentinel panel design. This came at the expense of significant spread into the BV711 channel (assigned to

PD-1). PD-1$^{hi}$ CD19$^{+}$ cells are rare in blood, for which this panel is designed, instead being predominantly found in tumour tissue (30).

- CD14 to cFluorB548, which has a unique spectrum and narrow primary emission peak, fully compatible with other fluorochromes with peak 488nm excitation. Permeabilisation changes cell morphology, making FSC/SSC an insufficient discriminator of the lymphocyte gate. Furthermore, monocytes have a unique autofluorescence spectrum, which could introduce unmixing errors. Lastly, monocytes degraded some of the tandem fluorochromes we tested, such as PE-Fire 810 (Supplementary Figure 5). This phenomenon has previously been reported for other APC and PE- tandem dyes(31). This necessitated a reliable monocyte exclusion marker on the panel, though it is not designed for the analysis of these cells.

##### 1.3. Selection of viability stain

Live Dead Blue viability dye was selected because its secondary emission peak is lower than that of other UV-excitable viability dyes, and it was successfully used in OMIP-069 (32).

##### 1.4. Allocation of secondary and tertiary markers

FOXP3, CD25 and HELIOS are tertiary markers necessary for accurate Treg identification and were therefore assigned first. Detection of FOXP3 can be enhanced by the combination of 2 antibody clones (33). In our laboratory, we routinely use the same principle for CD25. Therefore, they were assigned to PE and BV421 respectively - bright non-tandem dyes available conjugated to multiple antibody clones. HELIOS was assigned to PE-Dazzle 594, which is highly dissimilar to PE (SI 0.49). IgD was assigned to the highly similar Spark Yellow Green 581, as it is mutually exclusive with FOXP3- PE.

Antibodies for the chemokine receptors CCR4, CCR6 and CXCR3 were assigned early on based on successful prior use in our laboratory with cryopreserved samples. CXCR5 was added at the end of panel development, with Spark NIR 685 one of few remaining options. Since B-cells do not express CD127 (assigned to APCR700 as per Sahir et al (23), this allowed optimum CXCR5 detection on B-cells. CXCR5 expression on T-cells is lower, and due to spread from APC-CD27, we would recommend the use of an FMO control for CXCR5.

The remaining markers, which convey information about cell function, were allocated to fluorochromes with SI typically below 0.8, considering predicted co-expression patterns in healthy blood, as follows:

- IgM (BUV496) shares a fluorochrome with CD3 in a sentinel approach, as this fluorochrome was readily available and compatible with the rest of the panel. The only alternative at the time of panel design was conjugated to Alexa Fluor 350, which was not selected due to economic and availability reasons. Further options have been developed since, but their high similarity indices would require careful validation for compatibility with the panel, likely without any gains in resolution.

- CD21 (BUV615), is highly expressed by B-cells, and lower on naïve recent thymic emigrant T-effectors. It is not significantly expressed on Tregs, limiting any potential loss of resolution due to co-expression of FOXP3 (PE) and CD39 (BUV661).

- CD39 (BUV661), is expressed on B-cells, and on Tregs, where inter-individual variation is genetically determined(34). Some Tregs express CD49d (assigned BUV737 based on prior experience), but typically CD39 expression is high enough to avoid any loss of resolution.

- CD73 (BUV805), expressed on some B-cells, some CD4/CD8 T-effectors and very few Tregs, was chosen based on reagent availability.

- CD226 was assigned to BV510 due to high levels of expression on T- effector, B- , NK- cells and NKT cells. This fluorochrome limited the potential impact on other important markers on these populations. Although auto-fluorescence impacts this part of the spectrum, CD226 expression was still clearly detectable. The use of an FMO control aids gating on populations where it is dimmer, such as Tregs.

- HLA-DR (BV570) is also widely expressed on activated lymphocytes, at lower density relative to monocytes and B-cells. BV570 is a dim fluorochrome, which further limited any potential loss of resolution.

- PD-1 (BV711) was used in our previous panels. We would recommend an FMO control.

- CD24 (BV750) is not expressed significantly on T-cells, thereby limiting its impact on the detection of CD70 (BV786- limited by reagent availability) on T-cells. Antigen-activated B-cells express CD70, but CD24 expression does not negatively impact on its resolution. An FMO control is needed for both markers.

- TIGIT (PerCPefluor710) was compatible with all other 488nm-excited fluorochromes on the panel, sufficiently bright, and well resolved against CD226 (BV510), with which it competes for the same ligands.

- CD31 (BV480) is highly expressed on naïve T-cells, which conversely express little CD25 (BV421) and CD226 (BV510), making this a suitable combination.

##### 1.5. Allocation of remaining intracellular tertiary markers

TBET required a bright fluorochrome which is not impacted by other markers on the panel in all three lymphocyte lineages, making Kiravia Blue 520 (B2 channel) an optimal choice. For the same reason, this channel is customizable and can accommodate other markers, usually without extensive further validation.

CTLA-4 (PECy5) is highly sensitive to light degradation, so this fluorochrome was well suited to the final stages of staining, and sufficiently bright to avoid impact from CD27 (APC).

Initially, ki67 was assigned to PE-Cy5.5, expecting that proliferating cells would be rare, and minimise the potential impact of this fluorochrome with multiple emission peaks. However, its spread into BB700 despite down-titration (to 1:1000) prohibited its use. Finally, we assigned Ki-67 to efluor450, which was compatible with CD25 (BV421) and CD31 (BV480).

Finally, we had planned to include GATA3 to detect Th2 cells but detection was variable and poor (on APC). We replaced it with CRTH2, a surface marker more specific for Th2 lineage commitment (23).

Final panel optimisation required the relative comparison of CD3, CD4, CD8 and CD45RA density to reallocate CD45RA to BUV395 (originally planned for PerCP), and CD8 to PerCP (originally BUV496). Examples of unsuccessful early fluorochrome assignments are also shown (Supplementary Figure 6). The Spillover spreading matrix, calculated in FlowJo using unmixed single stain reference samples of cells stained with the optimised panel, is shown in Supplementary Figure 7.

##### 1.6. Antibody titration

All antibodies were titrated under identical staining conditions, using 2-fold serial dilutions between 1:25 and 1:1600 in 100μL staining volume. Dead cells were excluded with 1:500 Live Dead Blue (for fluorochromes emitting at long wavelengths) or Zombie NIR 1:3000 (for fluorochromes emitting in the UV and violet end of the spectrum). The number of total events collected was standardised across samples. Stain index analysis was performed on unmixed data. In addition to stain index, the frequency of the positive population, spread, total expected number of cells in the multi-stained samples (10-20-fold higher) were also considered when selecting optimum concentrations (Supplementary Figure 8). Optimum antibody amounts are shown in Supplementary Table 4.

#### 2. Supplementary Information- Panel testing

##### 2.1. Reference control optimisation

Multi-stained samples were unmixed with UltraComp bead controls (1:100 dilution of antibody) or cells stained at the optimum selected concentration based on titration. Bead- and cell- normalised unmixed reference control spectra were compared for discrepancies (Supplementary Figure 9). Cellular controls were preferred where the two methods were equivalent.

Bead controls were not suitable for BV786 unmixing. Due to the low expression of CD70 (BV786) on healthy control PBMCs, *in vitro* stimulated cells were used as an unmixing control. Specifically, we used cryopreserved stock $CD127^{lo}CD25^{hi}$ cells expanded for >4 rounds with CD3/28 beads and IL-2 according to our previously published protocol (35), as they express abundant CD70 and we had banked stocks. Alternatively, *in vitro* activated T-cells and certain human haematopoietic cell lines also express this marker.

##### 2.2. Unmixing optimisation

In cryopreserved PBMCs, auto-fluorescence (AF) was present in the UV 520-550nm and Violet 520-575nm region of the spectrum, corresponding to UV7-9 and V7 detectors on the Cytek Aurora cytometer (Supplementary Figure 10, a). This was not fully addressed by using Automatic AF extraction in Spectroflo v3.0. Therefore, other unmixing methods were compared by inspecting scatter plots of the affected channels. The optimal method involved exporting the 514nm+ (channel UV7) events from the unstained control of each experiment as a new FCS file, then importing them as a new fluorochrome reference control. The negative population for this control was the lymphocyte gate of the total unstained sample. Unmixing in updated versions of Spectroflo software, or in other spectral cytometry platforms, has not been tested and may require a different approach due to differences in the extraction algorithm.

##### 2.3. Unmixing accuracy in the multi-stained samples

Data were unmixed in Spectroflo v3.0, which uses an ordinary least squares algorithm, then cleaned of debri, aggregates, doubles, dead cells and CD14+ monocytes. The multi-stained samples and reference controls were then visualised in NxN plot format to determine unmixing accuracy. Spillover corrections were introduced where necessary by visually aligning the medians of the positive and negative populations, based on the reference controls and on prior knowledge of the expected expression patterns of the markers. Of 1156 combinations (excluding the autofluorescence parameters), 24 required spillover correction at or greater than +/-1%, and not exceeding 7% (between BUV496 and BUV563). An example of corrected and uncorrected data for two of the fluorochromes requiring the most significant correction is shown in Supplementary Figure 11.

##### 2.4. Evaluation of the resolution of each marker using single stain vs multi-stain comparisons

Adequate marker resolution, at the optimum antibody titre was evaluated by comparing the shape of the distribution of fluorescence signal in single stained- and multi-stained samples from the same donor (Supplementary Figure 12). The width of the negative population was similar in all samples and all markers, indicating that the effect of spread on resolution was likely minimal.

FMO controls were used to confirm resolution of each marker on key cell populations, and to aid gating (Supplementary Figure 13).

Finally, visual inspection of NxN plots determined that resolution of markers stained with highly similar fluorochromes was preserved in multi-stained samples (Supplementary Figure 14).

Healthy control donor cells have natural heterogeneity. For example, the donor used for the preparation of reference controls and FMOs expressed no TBET on their Th1 T-effectors (and only on CD8+ and NK cells). Selected examples of marker expression variability in healthy donors are shown in Supplementary Figure 15.

### 3. Supplementary information- Protocol and Methods

#### 3.1. Required materials for staining procedure:

- 15ml and 50ml Conical centrifuge tubes (Falcon): catalogue #E1415-0200 & E1450-0200
- 12x75 mm (5 ml) FACS tubes with cell strainer (Falcon®): catalogue # 352235
- 96 V-bottom well plates (Elkay): catalogue # MICR-TPV
- Brown 1.5 ml Safeseal tubes (Sarstedt): catalogue #72.706.001
- Cell Freezing medium (Merck): catalogue #C6164
- X-VIVO15 (Lonza): catalogue #02-060F
- PBS (Merck) catalogue #P2272-500ML
- Human AB Serum (Merck): catalogue # H4522-100ml
- Mouse serum (Merck) catalogue #M5905-10ML, heat inactivated for 20min at 56°C
- Fetal bovine serum (Merck): catalogue #F7524-500ML, heat inactivated as above
- FcR Blocker (Miltenyi) catalogue #130-059-901
- Penicillin-Streptomycin (Merck) catalogue #P0781-100ml
- Brilliant Staining buffer Plus (BD Biosciences), catalogue #566385
- True-Stain Monocyte Blocker™ (BioLegend) catalogue #426103
- CellBlox Blocking Buffer (ThermoFisher): catalogue #B001T03F01 (for use with Novafluor dyes)
- FoxP3 eBioScience staining buffer kit (ThermoFisher) catalogue #00-5523-00
- UltraComp eBeads™ Compensation Beads (Thermo Fisher): catalogue #01-2222-42
- Antibodies specified in Supplementary Table 4.

**Prepared buffers:**

- FACS buffer: PBS, 3% Human AB serum, 1% Penicillin-Streptomycin
- Thawing medium: X-VIVO15, 1% penicillin-streptomycin, 10% FBS

• Cell resting medium: X-VIVO15, 1% penicillin-streptomycin, 3% human AB serum

##### 3.2. Preparation of PBMCs

Blood was collected by peripheral venepuncture in sodium heparin tubes and processed after 20min rest at room temperature. PBMCs were isolated by density gradient centrifugation over Ficoll Paque Plus (GE Healthcare) according to the manufacturer's instructions. Briefly, blood was diluted 1:1 in PBS, layered over 15ml Ficoll Paque Plus in 50ml Falcon tubes and centrifuged at 400G for 25min with the brake off. The PBMC layer was aspirated from the buffy coat with a Pasteur pipette and washed twice in 50ml PBS, then resuspended in ice-cold Cell Freezing Medium (Sigma) by dropwise addition at 10 x $10^6$ cells/ml, followed by cryopreservation in cryovials (Star Lab) in CoolCell LX container overnight (-80°C) and long-term storage in liquid nitrogen.

##### 3.3. Thawing cells:

Cryogenic vials containing PBMCs were agitated in a 37°C water bath until almost completely thawed. 1ml pre-warmed X-VIVO15/10% FCS was added quickly to the vial. The suspension was transferred into a 15ml or 50ml Falcon tube containing the same medium, at a 1:10 dilution, centrifuged at 450G for 10min, re-suspended in X-VIVO15/3% human AB serum and rested at room temperature for 1 hour while preparing antibody master mixes.

##### 3.4. Staining procedure

All steps are carried out protected from light. Centrifugation and wash steps are always at the temperature specified in the preceding incubation step (i.e. either 4°C or 21°C), unless specified.

1. Thaw PBMCs. Pellet and resuspended in FACS buffer (3% human AB serum, 1% penicillin-streoptomycin, PBS), 5µL/test of FcR block, and 5µL/test of mouse serum, to a total volume of 20µL/sample plus 10% excess.

2. Add 20µL of cell suspension to brown Safeseal tubes containing the single chemokine receptor antibody or its master mix (with 5µL BD Brilliant Stain buffer Plus, 5µL of TrueStain Monocyte block and 10µL FACS buffer). Total volume is 50µL/test including cells. Incubate at room temperature for 15min.

3. While the brown tubes are incubating, add 20µL cell suspension to all single stain reference control wells (96 V-bottom well plate) on ice. Each reference well has a total staining volume of 100µL and contains TrueStain monocyte block (for CD4-NovaFluor610/70S fluorochrome, and all multi-stained samples, also add the manufacturer-recommended CellBlox buffer).

4. At the end of 15 min, transfer the chemokine receptor cell mix from the brown tubes to their corresponding position in the 96-well plate (4°C) and mix well.

5. Continue incubating all wells for 45min (4°C).

6. Wash twice in PBS.

7. Perform Viability staining with Live Dead Blue fixable viability dye (ThermoFisher). For the single-stain reference control, spike in dead cells (to produce a bright uniform positive population, we recommend using cells permeabilised with cold methanol for 5min, then washed in PBS). Thaw aliquot of Live/Dead Blue (dried preparation is resuspended in DMSO following the manufacturer recommendation and 5µL aliquots are stored at -20°C). The stock viability dye is used at a final concentration of 1:500 per test in PBS and total volume is 180µL/well (i.e. 0.36µL/test of stock solution; pre-dilute the stock aliquots in a volume of N x 5µL where N is the number of wells being stained, use 5µL per sample). Stain for 20min at 4°C. Stop the reaction with a droplet of FACS buffer per well, centrifuge (600G, 5min) and wash twice with PBS. Allow the plate to briefly equilibrate to room temperature on the bench for the next step.

8. Fix/permeabilise all wells using eBioscience FOXP3 Transcription Factor buffer set according to manufacturer's instructions for precisely 25min at room temperature. At the end, add a droplet of cold eBiosciences Perm/wash buffer, incubate for 5min (4°C, protected from light) and centrifuge at 700G for 5min (4°C), and wash twice more in Perm/wash buffer.

9. Re-suspend cells in intracellular antibody master-mix (prepared in permeabilisation buffer) for 60min at 4°C (agitating at 30min).

10. Wash twice with cold permeabilisation buffer, once in cold PBS and re-suspend in PBS. Keep samples cold and protected from light, and acquire within 24 hours.

#### 4. Supplementary information- Data acquisition and analysis

Samples were acquired on a 5-laser Cytek Aurora Cytometer with Cytek Automatic Settings following daily QC (Cytek QC bead lot 2003). Data were unmixed in Spectroflo v3.0.

Single stained controls were used to make minor corrections to Spillover (Supplementary Figure 11). Data were then cleaned of debris, doublets, dead cells and monocytes. Cleaned compensated unmixed FCS files were exported, and re-imported into FlowJo v10.8.1 for further analysis.

For high-dimensional analysis in Cytobank or FCS Express v7, healthy donor files were concatenated, data were bi-exponentially transformed and all channels were visually inspected to ensure population distributions were not distorted. Live CD14- single cells were subjected to dimensionality reduction using tSNE-CUDA or opt-SNE algorithm. FlowSOM unsupervised clustering analysis was performed on total T, B and NK cells with the settings specified in figure legends. Heatmaps were used to confirm the identity of the resulting populations and where necessary, clusters were manually separated or merged.

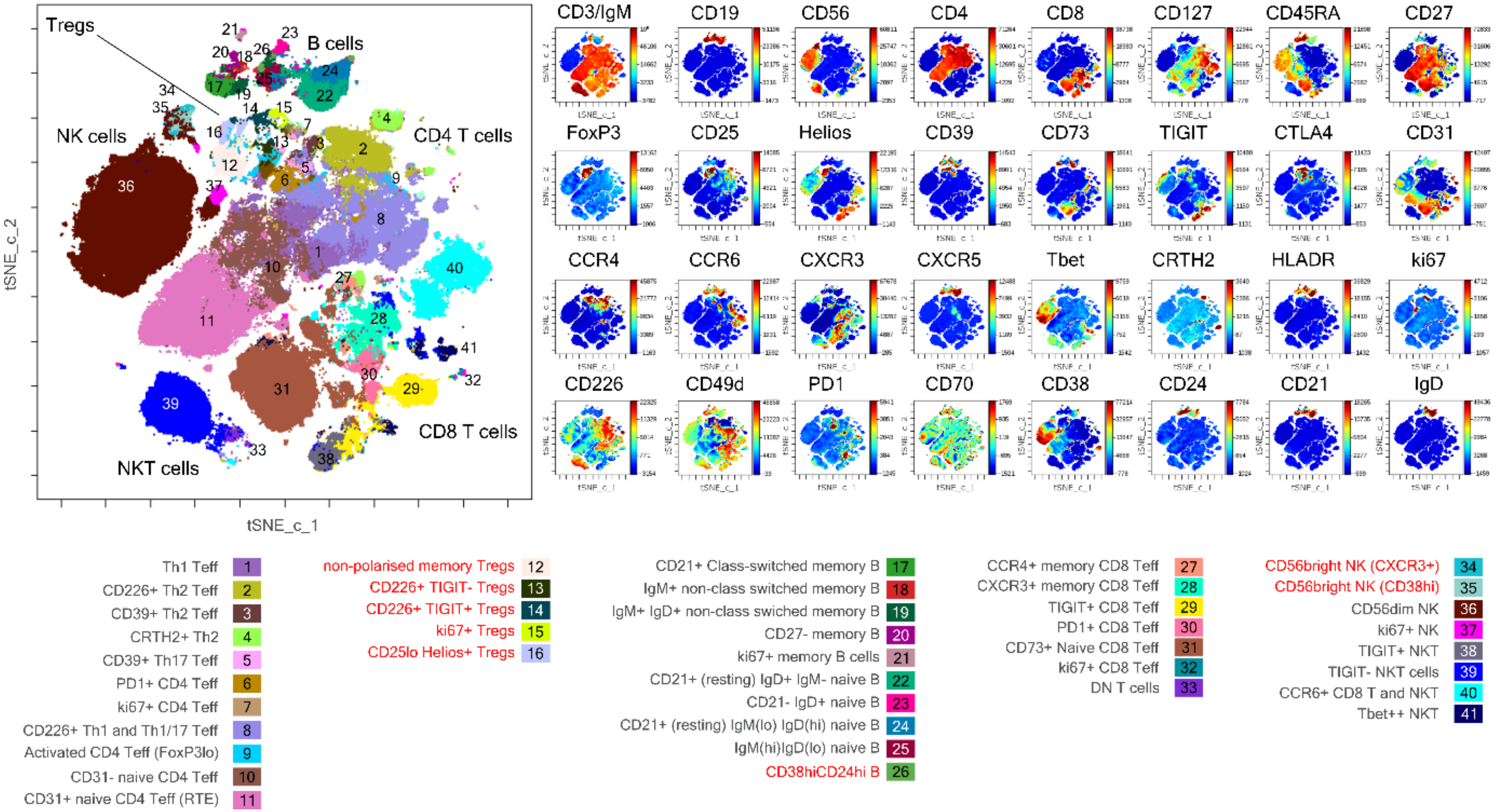


*Supplementary Figure 1 Dimensionality reduced analysis of 6 concatenated healthy donor samples by FlowSOM-on-tSNE on live single CD14- cells in Cytobank. 500,000 events were proportionally downsampled from the file. Scales were normalised and all 32 fluorescence channels were used for clustering (excluding viability and CD14) to perform consensus clustering with 45 metaclusters (400 clusters), seed 5875. Clusters were manually annotated based on marker expression by examining heatmaps and dot plots, and where necessary, they were merged or separated. Lineage-negative metaclusters, which likely represent dendritic cells and other non-lymphoid cells, have been excluded from the visualisation, as they could not be accurately identified with these markers.*

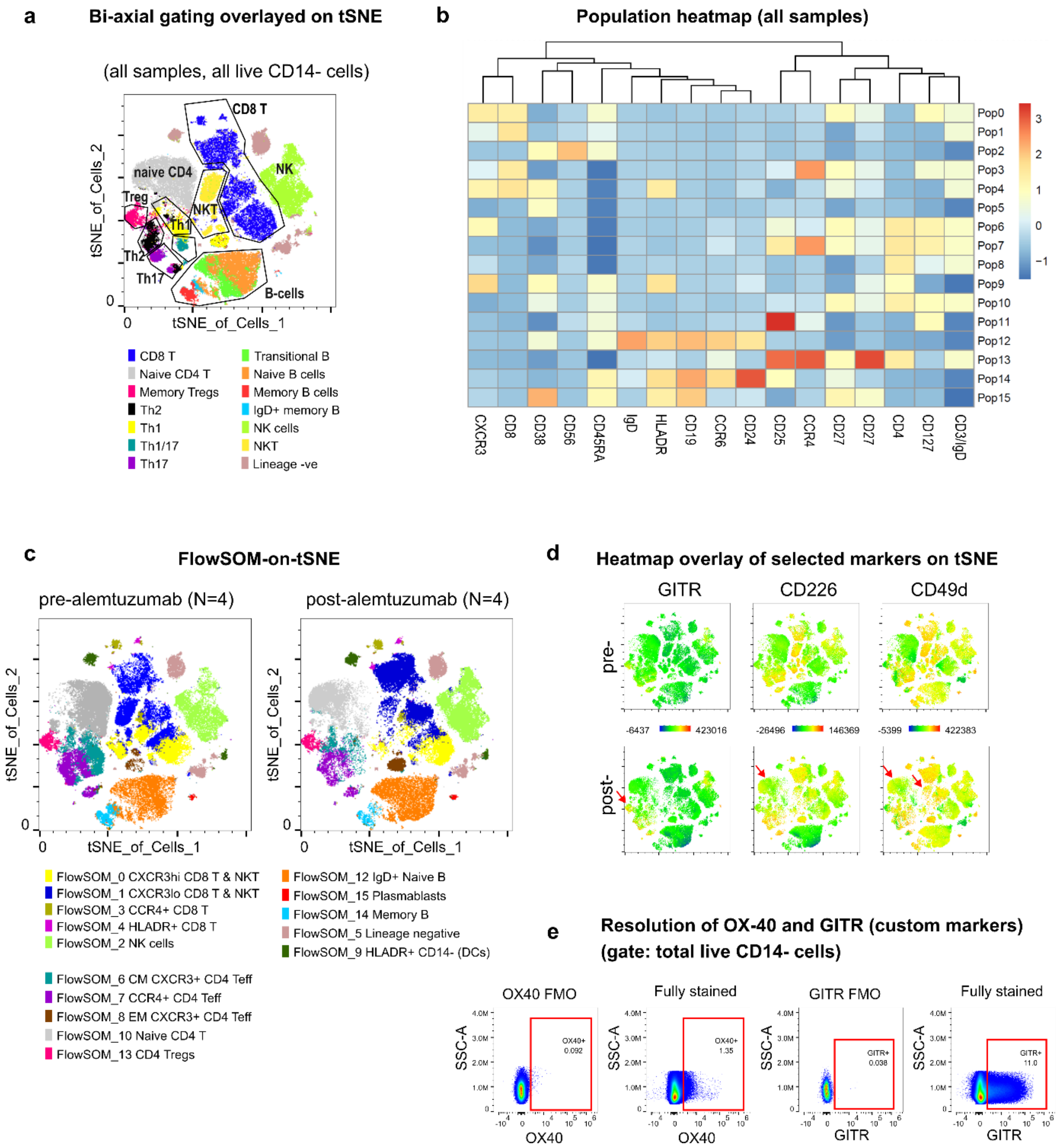


*Supplementary Figure 2 Customised version of the OMIP panel. TBET and CRTH2 were replaced with GITR and OX-40 respectively. PBMCs from 4 donors pre- and post- alemtuzumab treatment for multiple sclerosis were stained. 40,000 events per file were randomly downsampled (Downsamplev3, FlowJo), concatenated, exported as new FCS files and subjected to opt-SNE in FlowJo v10.8.1 (perplexity= 1000; eta= 30, KNN method= ANNOY, Barnes Hut approximation) using the following parameters: CD3/IgM, CD4, CD8, CD127, CD45RA, CD27, CD19, CD56, IgD, CD24, CD38, FoxP3, HLADR, CXCR3, CCR4, CCR6; followed by FlowSOM using the same markers (grid 16 x 16, metaclusters =16, seed= 42, with normalisation). A, Overlay of conventionally gated populations on dimensionality reduced map. B, Heatmap of marker expression by metacluster. C, Comparison of pre- and post-alemtuzumab T, B and NK lymphocytes coloured by FlowSOM populations showing changes in the topography of the naïve CD4, Treg and NK cell populations. D, Heatmap overlay of selected markers on tSNE. E, Manual gating showing the gate boundary for the custom markers in this version of the panel. Red arrows- post alemtuzumab, naïve CD4 T-cells gain expressed of CD226 and CD49d; Tregs and CD8/NKT cells also undergo changes. These changes have also been reported by others ((36)), using multiple small panels which requires more cellular material.*

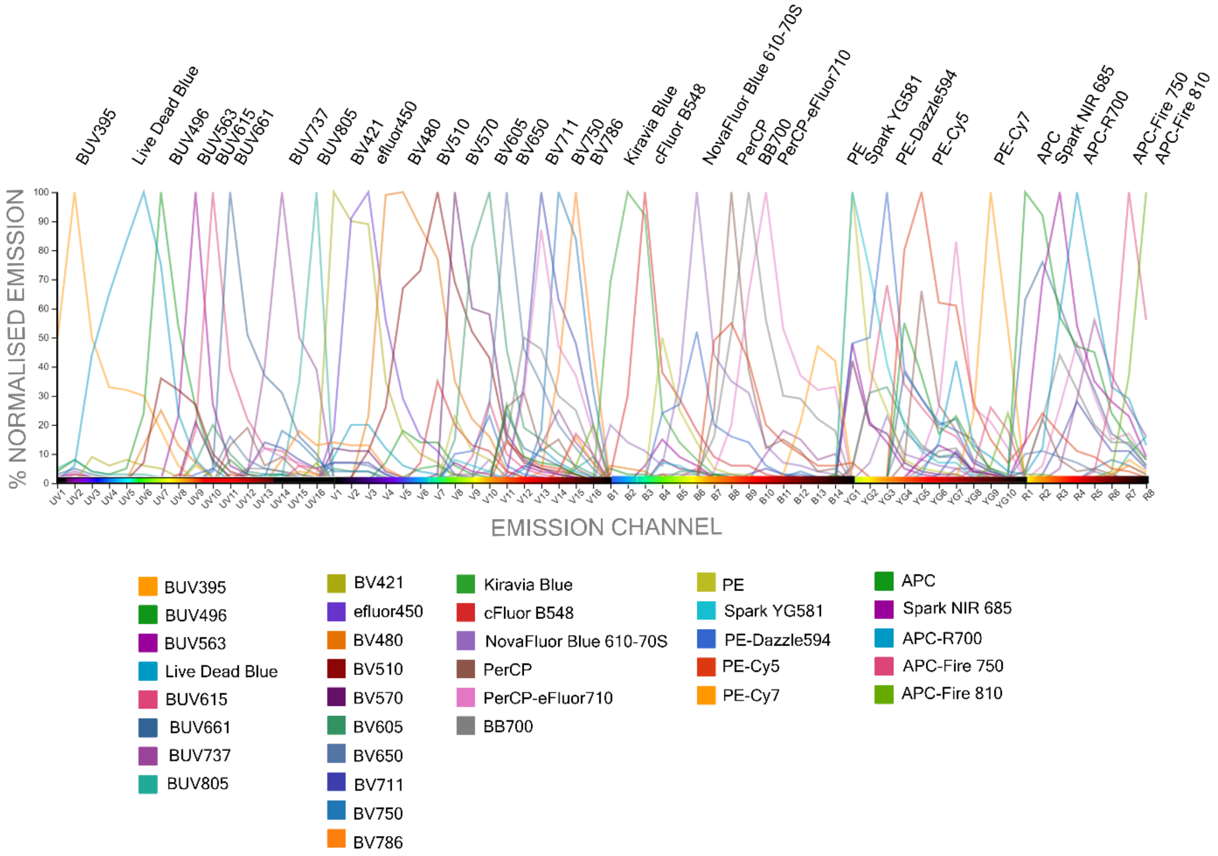


*Supplementary Figure 3 Published normalised spectra of the fluorochromes used in this OMIP. Source- [https://cloud.cytekbio.com/newlogin](https://cloud.cytekbio.com/newlogin) CytekBio Spectral viewer.*

#### Similarity™ Indices

**Configuration 5L 16UV-14B-10YG-8R**

| | BUV395 | BUV496 | BUV563 | LIVE DEAD Blue | BUV615 | BUV661 | BUV737 | BUV805 | BV421 | eFluor 450 | BV480 | BV510 | BV570 | BV605 | BV650 | BV711 | BV750 | BV786 | KIRAVIA Blue 520 | cFluor B548 | NovaFluor Blue 610-70S | PerCP | PerCP-eFluor 710 | BB700 | PE | Spark YG 581 | PE-Dazzle594 | PE-Cy5 | PE-Cy7 | APC | Spark NIR 685 | APC-R700 | APC-Fire 750 | APC-Fire 810 |
|---|---|---|---|---|---|---|---|---|---|---|---|---|---|---|---|---|---|---|---|---|---|---|---|---|---|---|---|---|---|---|---|---|---|---|
| BUV395 | 1 | | | | | | | | | | | | | | | | | | | | | | | | | | | | | | | | | |
| BUV496 | 0.25 | 1 | | | | | | | | | | | | | | | | | | | | | | | | | | | | | | | | |
| BUV563 | 0.05 | 0.29 | 1 | | | | | | | | | | | | | | | | | | | | | | | | | | | | | | | |
| LIVE DEAD Blue | 0.51 | 0.58 | 0.09 | 1 | | | | | | | | | | | | | | | | | | | | | | | | | | | | | | |
| BUV615 | 0.03 | 0.08 | 0.4 | 0.03 | 1 | | | | | | | | | | | | | | | | | | | | | | | | | | | | | |
| BUV661 | 0.04 | 0.02 | 0.06 | 0.03 | 0.35 | 1 | | | | | | | | | | | | | | | | | | | | | | | | | | | | |
| BUV737 | 0.03 | 0.01 | 0.01 | 0.02 | 0.1 | 0.37 | 1 | | | | | | | | | | | | | | | | | | | | | | | | | | | |
| BUV805 | 0.11 | 0.03 | 0.01 | 0.06 | 0.04 | 0.1 | 0.39 | 1 | | | | | | | | | | | | | | | | | | | | | | | | | | |
| BV421 | 0.05 | 0.09 | 0.01 | 0.26 | 0 | 0 | 0 | 0.01 | 1 | | | | | | | | | | | | | | | | | | | | | | | | | |
| eFluor 450 | 0 | 0.08 | 0 | 0.21 | 0 | 0 | 0 | 0 | 0.85 | 1 | | | | | | | | | | | | | | | | | | | | | | | | |
| BV480 | 0.02 | 0.38 | 0.07 | 0.2 | 0.04 | 0.01 | 0 | 0 | 0.28 | 0.46 | 1 | | | | | | | | | | | | | | | | | | | | | | | |
| BV510 | 0.02 | 0.48 | 0.22 | 0.2 | 0.14 | 0.04 | 0.02 | 0 | 0.17 | 0.27 | 0.84 | 1 | | | | | | | | | | | | | | | | | | | | | | |
| BV570 | 0.01 | 0.09 | 0.37 | 0.04 | 0.29 | 0.06 | 0.02 | 0 | 0.15 | 0.14 | 0.26 | 0.54 | 1 | | | | | | | | | | | | | | | | | | | | | |
| BV605 | 0.01 | 0.05 | 0.2 | 0.02 | 0.51 | 0.17 | 0.05 | 0.01 | 0.06 | 0.06 | 0.16 | 0.4 | 0.7 | 1 | | | | | | | | | | | | | | | | | | | | |
| BV650 | 0 | 0.02 | 0.04 | 0.02 | 0.26 | 0.4 | 0.13 | 0.02 | 0.09 | 0.08 | 0.06 | 0.16 | 0.25 | 0.53 | 1 | | | | | | | | | | | | | | | | | | | |
| BV711 | 0 | 0 | 0.01 | 0.02 | 0.07 | 0.25 | 0.41 | 0.09 | 0.09 | 0.08 | 0.03 | 0.06 | 0.07 | 0.17 | 0.45 | 1 | | | | | | | | | | | | | | | | | | |
| BV750 | 0 | 0 | 0 | 0.01 | 0.03 | 0.09 | 0.37 | 0.14 | 0.05 | 0.04 | 0.02 | 0.04 | 0.04 | 0.1 | 0.23 | 0.69 | 1 | | | | | | | | | | | | | | | | | |
| BV786 | 0 | 0.01 | 0 | 0.04 | 0.03 | 0.07 | 0.29 | 0.23 | 0.2 | 0.17 | 0.05 | 0.06 | 0.06 | 0.09 | 0.19 | 0.54 | 0.86 | 1 | | | | | | | | | | | | | | | | |
| KIRAVIA Blue 520 | 0.01 | 0.07 | 0.06 | 0.01 | 0.01 | 0 | 0 | 0 | 0.01 | 0.01 | 0.09 | 0.05 | 0.03 | 0.01 | 0 | 0 | 0 | 0 | 1 | | | | | | | | | | | | | | | |
| cFluor B548 | 0.02 | 0.09 | 0.14 | 0.01 | 0.04 | 0.01 | 0.01 | 0 | 0.02 | 0.03 | 0.21 | 0.3 | 0.24 | 0.16 | 0.06 | 0.02 | 0.01 | 0.02 | 0.73 | 1 | | | | | | | | | | | | | | |
| NovaFluor Blue 610-70S | 0.01 | 0.01 | 0.12 | 0 | 0.19 | 0.05 | 0.02 | 0 | 0 | 0 | 0.02 | 0.02 | 0.11 | 0.17 | 0.06 | 0.02 | 0 | 0 | 0.27 | 0.34 | 1 | | | | | | | | | | | | | |
| PerCP | 0 | 0.01 | 0.04 | 0.01 | 0.22 | 0.38 | 0.15 | 0.03 | 0.01 | 0.01 | 0.01 | 0.01 | 0.06 | 0.18 | 0.38 | 0.23 | 0.13 | 0.08 | 0 | 0.06 | 0.36 | 1 | | | | | | | | | | | | |
| PerCP-eFluor 710 | 0 | 0 | 0.02 | 0 | 0.12 | 0.29 | 0.39 | 0.07 | 0 | 0 | 0 | 0.02 | 0.05 | 0.14 | 0.31 | 0.65 | 0.38 | 0.28 | 0.01 | 0.05 | 0.21 | 0.46 | 1 | | | | | | | | | | | |
| BB700 | 0 | 0 | 0.02 | 0 | 0.08 | 0.33 | 0.3 | 0.05 | 0 | 0 | 0.02 | 0.05 | 0.08 | 0.16 | 0.37 | 0.53 | 0.3 | 0.22 | 0.03 | 0.11 | 0.34 | 0.64 | 0.82 | 1 | | | | | | | | | | |
| PE | 0.01 | 0.03 | 0.51 | 0 | 0.25 | 0.03 | 0 | 0 | 0 | 0.01 | 0.05 | 0.12 | 0.52 | 0.28 | 0.05 | 0.01 | 0 | 0 | 0.09 | 0.24 | 0.25 | 0.06 | 0.03 | 0.06 | 1 | | | | | | | | | |
| Spark YG 581 | 0 | 0.01 | 0.46 | 0 | 0.36 | 0.06 | 0 | 0 | 0 | 0 | 0.02 | 0.04 | 0.42 | 0.3 | 0.06 | 0.01 | 0 | 0 | 0.02 | 0.05 | 0.15 | 0.07 | 0.04 | 0.03 | 0.88 | 1 | | | | | | | | |
| PE-Dazzle594 | 0.01 | 0.02 | 0.36 | 0 | 0.58 | 0.13 | 0.02 | 0 | 0 | 0 | 0.04 | 0.09 | 0.41 | 0.47 | 0.16 | 0.04 | 0.01 | 0.01 | 0.07 | 0.2 | 0.6 | 0.27 | 0.16 | 0.18 | 0.67 | 0.72 | 1 | | | | | | | |
| PE-Cy5 | 0 | 0 | 0.07 | 0 | 0.32 | 0.42 | 0.1 | 0.01 | 0 | 0 | 0 | 0.01 | 0.09 | 0.21 | 0.29 | 0.14 | 0.05 | 0.03 | 0.01 | 0.06 | 0.35 | 0.77 | 0.47 | 0.48 | 0.14 | 0.19 | 0.43 | 1 | | | | | | |
| PE-Cy7 | 0 | 0 | 0.01 | 0 | 0.04 | 0.05 | 0.14 | 0.08 | 0 | 0 | 0 | 0 | 0.01 | 0.03 | 0.04 | 0.11 | 0.15 | 0.17 | 0 | 0.01 | 0.04 | 0.09 | 0.29 | 0.17 | 0.02 | 0.03 | 0.05 | 0.14 | 1 | | | | | |
| APC | 0 | 0 | 0.03 | 0.01 | 0.17 | 0.78 | 0.24 | 0.02 | 0 | 0 | 0 | 0.02 | 0.05 | 0.14 | 0.37 | 0.23 | 0.07 | 0.04 | 0 | 0.01 | 0.06 | 0.34 | 0.29 | 0.33 | 0.04 | 0.08 | 0.16 | 0.52 | 0.05 | 1 | | | | |
| Spark NIR 685 | 0 | 0 | 0.01 | 0.01 | 0.05 | 0.65 | 0.28 | 0.03 | 0 | 0 | 0 | 0 | 0.01 | 0.03 | 0.19 | 0.27 | 0.06 | 0.04 | 0 | 0 | 0.01 | 0.21 | 0.31 | 0.41 | 0.01 | 0.02 | 0.04 | 0.28 | 0.04 | 0.75 | 1 | | | |
| APC-R700 | 0 | 0 | 0.01 | 0.01 | 0.06 | 0.5 | 0.45 | 0.04 | 0 | 0 | 0 | 0.01 | 0.02 | 0.05 | 0.18 | 0.39 | 0.11 | 0.07 | 0 | 0 | 0.02 | 0.17 | 0.49 | 0.41 | 0.01 | 0.02 | 0.06 | 0.28 | 0.1 | 0.57 | 0.78 | 1 | | |
| APC-Fire 750 | 0 | 0.01 | 0 | 0 | 0.02 | 0.13 | 0.31 | 0.21 | 0 | 0 | 0.01 | 0.01 | 0 | 0.02 | 0.05 | 0.19 | 0.2 | 0.23 | 0 | 0 | 0.01 | 0.04 | 0.18 | 0.15 | 0 | 0 | 0.01 | 0.07 | 0.32 | 0.16 | 0.23 | 0.3 | 1 | |
| APC-Fire 810 | 0 | 0 | 0 | 0 | 0.01 | 0.1 | 0.21 | 0.24 | 0 | 0 | 0 | 0 | 0 | 0.01 | 0.05 | 0.12 | 0.12 | 0.17 | 0 | 0 | 0 | 0.03 | 0.13 | 0.1 | 0 | 0 | 0.01 | 0.06 | 0.2 | 0.13 | 0.16 | 0.2 | 0.74 | 1 |

Color scale: 0 – 0.5 – 1.0

**Complexity™ index: 12.44**

*Supplementary Figure 4 Published similarity indices of the fluorochromes used in this OMIP, and overall complexity index of the panel.*

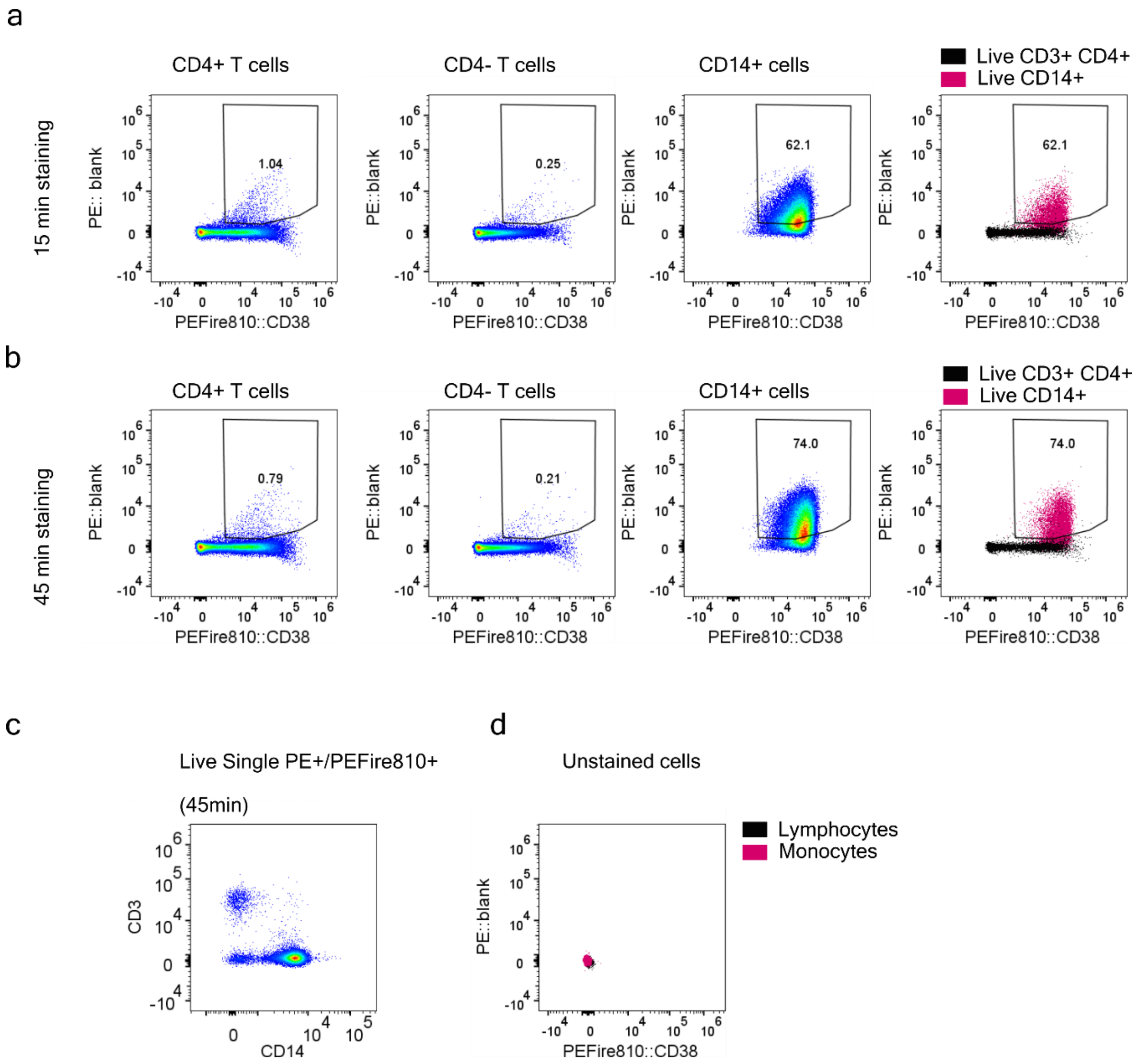


*Supplementary Figure 5 Monocytes degrade some tandem fluorochromes and must be excluded from analysis. Cryopreserved healthy donor PBMCs were surface- stained with the indicated fluorochromes for the indicated amount of time (a and b) on ice, protected from light, then fixed with 0.25% PFA for 15min and acquired. Multi-stained samples do not contain PE or APC. A, B- PE or APC signal in the multi-stained samples is due to fluorochrome degradation, highest on CD14+ cells and increasing with incubation time. C, the PE+ population consists mainly of monocytes. D, Unstained samples do not contain signal in PE (i.e. this is not an unmixing artefact).*

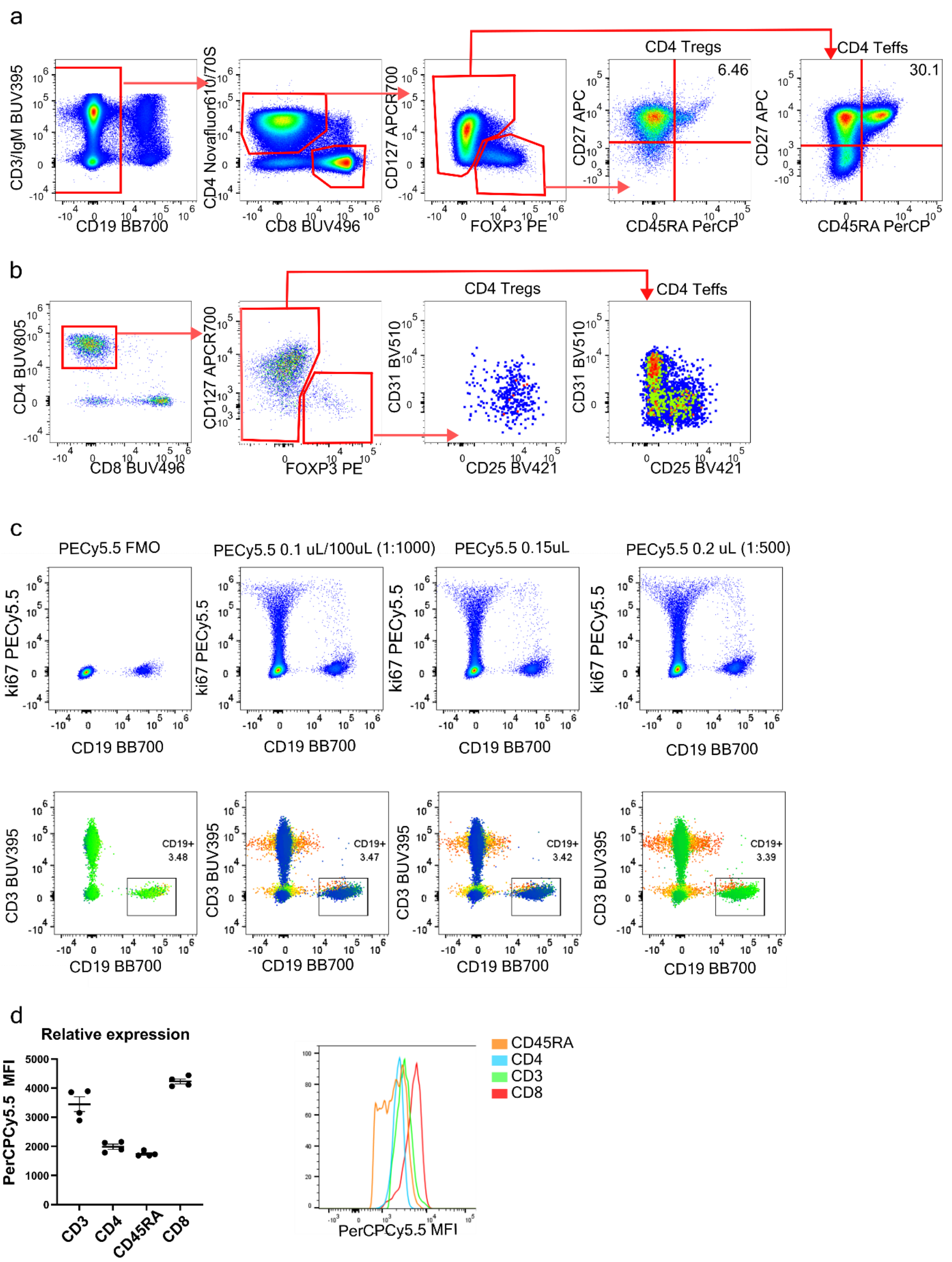


*Supplementary Figure 6 Examples of suboptimal fluorochrome assignments during panel design. A- CD45RA- PerCP resulted in poor resolution of naïve Tregs due to spread from multiple fluorochromes, including CD27-APC. B- CD31-BV510 was too dim to resolve well, especially on CD25+ naïve cells which arise during homeostatic turnover. C- ki67 on PECy5.5 spread into CD19, making separation of B-cells impossible; heatmap overlay= ki67 MFI. D- To reassign fluorochromes for CD8, CD45RA and CD3, their relative expression was determined on 4 healthy donors; all markers were conjugated to PerCPCy5.5 and stained in one batch. Representative histograms from one donor are shown.*

| | BUV395 | LDBlue | BUV496 | BUV563 | BUV615 | BUV661 | BUV737 | BUV805 | BV421 | ef450 | BV480 | BV510 | BV570 | BV605 | BV650 | BV711 | BV750 | BV786 | KB520 | cFB548 | NF610 70S | PerCP | BB700 | PCP710 | PE | SYG581 | PE-Daz | PE-Cy5 | PE-Cy7 | APC | SNIR685 | APCR700 | APC750 | APC810 | Sum |
|---|---|---|---|---|---|---|---|---|---|---|---|---|---|---|---|---|---|---|---|---|---|---|---|---|---|---|---|---|---|---|---|---|---|---|---|
| BUV395 | | 11.77 | 13.07 | 6.27 | 0.69 | 0.34 | 0.18 | 0.00 | 0.64 | 0.82 | 0.91 | 4.86 | 1.50 | 1.32 | 1.10 | 0.78 | 0.52 | 0.64 | 1.43 | 1.73 | 1.18 | 0.29 | 0.32 | 0.00 | 0.73 | 1.55 | 0.59 | 0.20 | 0.35 | 0.21 | 0.00 | 0.00 | 0.00 | 0.00 | 53.99 |
| LDBlue | 2.25 | | 12.20 | 5.447 | 0.89 | 0.61 | 0.32 | 0.00 | 1.10 | 2.04 | 1.44 | 4.005 | 1.31 | 1.39 | 1.20 | 1.07 | 0.64 | 0.83 | 1.15 | 1.19 | 0.86 | 0.48 | 0.58 | 0.30 | 1.61 | 1.30 | 0.91 | 0.01 | 0.24 | 0.47 | 0.32 | 0.46 | 0.37 | 0.44 | 47.43 |
| BUV496 | 1.07 | 1.36 | | 3.816 | 0.90 | 0.55 | 0.00 | 0.62 | 0.29 | 0.00 | 0.94 | 3.936 | 1.11 | 1.01 | 0.99 | 0.90 | 0.67 | 0.89 | 1.38 | 1.63 | 0.00 | 0.30 | 0.56 | 0.52 | 1.08 | 0.84 | 0.98 | 0.28 | 0.34 | 0.29 | 0.00 | 0.29 | 0.00 | 0.43 | 27.96 |
| BUV563 | 1.37 | 1.84 | 9.11 | | 1.50 | 0.93 | 0.75 | 0.61 | 0.60 | 1.20 | 0.77 | 6.345 | 1.76 | 0.73 | 0.00 | 0.00 | 0.00 | 0.00 | 2.04 | 2.42 | 1.86 | 0.64 | 0.69 | 0.73 | 4.253 | 2.20 | 1.63 | 0.75 | 0.01 | 0.44 | 0.75 | 0.44 | 0.71 | 0.64 | 47.71 |
| BUV615 | 0.82 | 0.00 | 0.00 | 0.88 | | 1.67 | 1.92 | 1.23 | 0.00 | 1.16 | 1.39 | 0.00 | 0.69 | 1.86 | 0.00 | 1.09 | 0.00 | 0.65 | 1.46 | 0.00 | 2.44 | 1.57 | 0.00 | 0.00 | 2.61 | 0.00 | 3.993 | 1.49 | 0.00 | 1.75 | 0.00 | 0.00 | 0.00 | 0.00 | 28.69 |
| BUV661 | 0.97 | 1.16 | 0.00 | 0.00 | 0.93 | | 2.44 | 2.00 | 0.00 | 0.96 | 0.00 | 3.462 | 0.06 | 1.21 | 2.26 | 0.92 | 0.00 | 1.24 | 1.05 | 0.00 | 0.00 | 2.03 | 1.53 | 1.31 | 1.17 | 1.93 | 1.47 | 2.46 | 0.00 | 3.566 | 2.98 | 2.06 | 1.39 | 0.85 | 41.39 |
| BUV737 | 0.41 | 0.77 | 0.00 | 0.00 | 0.00 | 0.77 | | 3.733 | 0.00 | 0.97 | 0.97 | 1.20 | 0.00 | 0.80 | 0.43 | 0.88 | 1.83 | 1.96 | 1.03 | 0.57 | 0.00 | 0.83 | 0.92 | 0.98 | 0.00 | 0.49 | 0.47 | 0.01 | 0.82 | 1.02 | 1.04 | 2.08 | 2.08 | 1.57 | 28.64 |
| BUV805 | 0.86 | 0.00 | 5.187 | 2.42 | 0.00 | 0.78 | 0.91 | | 0.02 | 0.99 | 1.12 | 2.29 | 1.25 | 2.18 | 3.043 | 2.42 | 0.68 | 1.53 | 2.93 | 0.81 | 0.71 | 0.86 | 1.35 | 0.00 | 1.70 | 0.08 | 0.98 | 0.80 | 1.12 | 0.00 | 0.91 | 1.01 | 1.53 | 1.39 | 41.86 |
| BV421 | 0.00 | 0.90 | 6.407 | 1.04 | 0.00 | 0.00 | 0.00 | 0.00 | | 2.95 | 1.16 | 0.00 | 0.57 | 0.00 | 0.05 | 0.74 | 1.46 | 0.00 | 1.22 | 1.48 | 0.98 | 0.00 | 0.00 | 0.97 | 0.00 | 0.83 | 0.00 | 0.00 | 0.56 | 0.47 | 0.00 | 0.00 | 0.83 | 0.00 | 22.61 |
| ef450 | 1.08 | 2.15 | 3.152 | 2.43 | 1.85 | 1.57 | 0.00 | 0.83 | 3.874 | | 3.639 | 9.98 | 3.713 | 0.68 | 0.00 | 0.00 | 2.25 | 0.00 | 1.05 | 3.061 | 2.48 | 0.00 | 1.19 | 0.00 | 1.22 | 2.05 | 1.45 | 1.46 | 0.51 | 0.00 | 2.02 | 0.00 | 0.88 | 0.62 | 55.18 |
| BV480 | 0.90 | 1.22 | 6.556 | 2.22 | 0.00 | 0.35 | 0.00 | 0.00 | 0.48 | 1.15 | | 3.678 | 2.67 | 1.73 | 0.96 | 0.56 | 0.00 | 0.00 | 1.41 | 2.20 | 0.97 | 0.64 | 0.70 | 0.52 | 1.28 | 0.45 | 0.43 | 0.34 | 0.61 | 0.00 | 0.00 | 0.51 | 0.63 | 0.00 | 33.15 |
| BV510 | 1.29 | 1.48 | 9.84 | 3.817 | 0.71 | 0.69 | 1.00 | 0.00 | 0.70 | 1.67 | 1.14 | | 3.336 | 2.68 | 0.78 | 0.80 | 1.31 | 1.57 | 2.54 | 3.093 | 1.82 | 0.73 | 0.57 | 0.52 | 3.031 | 1.57 | 2.08 | 0.00 | 0.42 | 1.44 | 0.55 | 0.00 | 0.90 | 0.91 | 53.00 |
| BV570 | 0.00 | 1.36 | 2.92 | 3.384 | 0.92 | 0.44 | 0.58 | 0.00 | 1.63 | 1.94 | 1.01 | 5.522 | | 3.235 | 1.22 | 0.80 | 0.64 | 0.37 | 1.78 | 1.97 | 2.02 | 1.02 | 1.18 | 0.75 | 3.517 | 2.58 | 1.35 | 0.96 | 0.67 | 0.32 | 0.00 | 0.72 | 0.00 | 0.32 | 45.10 |
| BV605 | 0.49 | 0.82 | 2.79 | 1.56 | 1.47 | 1.06 | 0.82 | 0.66 | 0.73 | 1.07 | 0.37 | 5.062 | 1.88 | | 1.99 | 0.92 | 1.12 | 1.60 | 1.80 | 1.57 | 1.91 | 1.18 | 1.44 | 0.70 | 2.74 | 2.64 | 2.14 | 1.24 | 0.40 | 1.23 | 0.51 | 0.00 | 0.00 | 0.61 | 44.51 |
| BV650 | 0.00 | 0.00 | 5.232 | 1.05 | 0.00 | 1.49 | 1.00 | 0.67 | 0.67 | 1.32 | 0.00 | 3.382 | 0.59 | 0.80 | | 1.97 | 1.62 | 2.13 | 0.05 | 1.83 | 0.00 | 1.69 | 1.50 | 0.89 | 0.00 | 0.00 | 0.58 | 1.10 | 0.83 | 0.70 | 0.54 | 0.69 | 0.50 | 0.51 | 33.34 |
| BV711 | 1.49 | 2.09 | 0.00 | 0.00 | 0.83 | 1.42 | 1.68 | 1.63 | 0.03 | 1.73 | 2.07 | 8.364 | 0.00 | 0.00 | 0.00 | | 3.395 | 2.40 | 0.00 | 0.00 | 1.02 | 0.86 | 0.00 | 0.00 | 0.00 | 0.00 | 0.00 | 0.81 | 0.02 | 1.43 | 0.90 | 1.87 | 1.72 | 2.14 | 37.89 |
| BV750 | 0.87 | 0.00 | 0.00 | 0.00 | 0.00 | 1.75 | 2.88 | 1.44 | 0.02 | 1.22 | 0.65 | 0.00 | 1.45 | 0.00 | 0.63 | 1.87 | | 4.147 | 2.34 | 2.21 | 1.00 | 0.00 | 1.16 | 0.00 | 0.85 | 0.00 | 0.68 | 0.00 | 0.00 | 0.00 | 0.00 | 0.00 | 0.85 | 0.00 | 26.03 |
| BV786 | 0.00 | 0.22 | 0.00 | 0.00 | 0.43 | 0.19 | 1.10 | 2.96 | 0.55 | 0.91 | 0.00 | 0.00 | 0.58 | 0.31 | 0.53 | 0.74 | 2.27 | | 0.77 | 1.02 | 0.53 | 0.45 | 0.45 | 0.35 | 0.69 | 0.00 | 0.40 | 0.19 | 0.51 | 0.44 | 0.43 | 0.27 | 1.57 | 1.23 | 20.11 |
| KB520 | 1.71 | 0.68 | 2.21 | 2.32 | 0.61 | 0.59 | 0.94 | 1.03 | 0.00 | 1.65 | 1.52 | 10.20 | 1.51 | 1.00 | 0.00 | 0.00 | 0.00 | 0.00 | | 5.998 | 1.05 | 0.90 | 1.41 | 0.00 | 2.67 | 1.76 | 0.00 | 0.86 | 0.51 | 0.89 | 1.53 | 0.62 | 0.89 | 0.91 | 45.97 |
| cFB548 | 0.79 | 1.24 | 8.416 | 3.439 | 0.55 | 0.61 | 0.48 | 0.44 | 0.48 | 1.21 | 3.739 | 6.178 | 2.28 | 1.37 | 0.99 | 0.82 | 0.71 | 0.33 | 3.282 | | 2.15 | 0.76 | 1.09 | 0.69 | 2.82 | 1.83 | 1.04 | 0.65 | 0.24 | 0.77 | 0.59 | 0.40 | 0.47 | 0.30 | 51.17 |
| NF610 70S | 0.59 | 1.32 | 3.503 | 0.92 | 0.62 | 0.64 | 0.38 | 0.29 | 0.35 | 1.46 | 0.68 | 7.864 | 1.66 | 1.97 | 1.22 | 1.32 | 0.60 | 0.89 | 3.294 | 3.874 | | 1.85 | 2.78 | 1.67 | 2.47 | 1.67 | 3.477 | 1.77 | 0.63 | 0.75 | 1.17 | 0.80 | 0.59 | 0.01 | 53.10 |
| PerCP | 0.00 | 0.91 | 0.00 | 0.00 | 0.41 | 1.88 | 1.93 | 1.54 | 0.81 | 0.00 | 0.90 | 2.69 | 0.50 | 0.94 | 1.48 | 1.26 | 2.07 | 1.57 | 0.53 | 0.99 | 0.00 | | 3.13 | 2.01 | 0.59 | 0.91 | 0.99 | 1.79 | 0.80 | 1.43 | 2.01 | 1.67 | 0.96 | 0.61 | 37.29 |
| BB700 | 0.00 | 0.85 | 3.164 | 0.00 | 0.53 | 1.08 | 1.06 | 1.04 | 0.75 | 0.00 | 0.00 | 4.254 | 0.03 | 0.36 | 1.38 | 3.829 | 2.53 | 2.66 | 1.15 | 1.79 | 1.15 | 1.76 | | 3.144 | 1.37 | 1.04 | 0.00 | 1.73 | 0.97 | 1.12 | 2.89 | 2.20 | 1.81 | 0.87 | 46.50 |
| PCP710 | 0.90 | 1.69 | 7.532 | 3.055 | 0.00 | 0.02 | 1.38 | 1.98 | 0.00 | 0.00 | 2.00 | 7.91 | 1.07 | 0.04 | 0.00 | 4.212 | 2.80 | 2.01 | 2.75 | 2.50 | 2.03 | 2.10 | 4.461 | | 3.309 | 1.58 | 0.72 | 1.36 | 1.76 | 1.83 | 2.56 | 2.54 | 2.00 | 1.79 | 69.90 |
| PE | 0.60 | 1.24 | 5.134 | 2.05 | 0.41 | 0.00 | 0.64 | 0.00 | 0.91 | 1.33 | 0.00 | 9.66 | 2.77 | 1.38 | 1.02 | 1.05 | 0.84 | 0.00 | 2.95 | 3.00 | 2.67 | 0.60 | 1.47 | 0.61 | | 4.206 | 1.60 | 0.93 | 0.80 | 1.22 | 0.00 | 0.41 | 0.00 | 0.00 | 49.49 |
| SYG 581 | 0.30 | 1.53 | 2.88 | 1.58 | 0.41 | 0.28 | 0.52 | 0.00 | 0.01 | 1.12 | 0.72 | 7.624 | 2.04 | 1.03 | 0.87 | 0.00 | 0.34 | 0.61 | 2.10 | 2.82 | 1.68 | 1.36 | 1.08 | 0.69 | 4.064 | | 2.31 | 1.27 | 0.00 | 0.29 | 0.55 | 0.51 | 0.52 | 0.02 | 41.13 |
| PE-Daz | 0.00 | 0.78 | 2.80 | 0.00 | 1.29 | 0.67 | 0.00 | 0.00 | 0.66 | 1.54 | 1.21 | 4.501 | 0.00 | 1.01 | 0.00 | 0.00 | 0.00 | 0.00 | 1.87 | 1.55 | 3.375 | 2.18 | 2.18 | 1.70 | 3.136 | 2.25 | | 2.01 | 0.00 | 1.23 | 0.53 | 1.09 | 0.72 | 0.88 | 39.16 |
| PE-Cy5 | 1.09 | 0.00 | 4.751 | 3.615 | 0.00 | 1.48 | 0.96 | 1.04 | 0.84 | 1.95 | 0.00 | 3.269 | 2.03 | 0.06 | 1.18 | 0.69 | 0.06 | 0.00 | 1.80 | 1.20 | 1.06 | 4.465 | 3.528 | 3.698 | 2.38 | 2.82 | 2.13 | | 0.92 | 2.25 | 2.44 | 2.12 | 1.29 | 0.91 | 56.01 |
| PE-Cy7 | 0.56 | 0.52 | 0.00 | 0.72 | 0.38 | 0.45 | 0.60 | 1.36 | 0.37 | 1.05 | 0.73 | 3.126 | 0.84 | 0.00 | 0.42 | 0.88 | 0.97 | 2.65 | 1.20 | 1.14 | 1.06 | 0.01 | 1.06 | 1.48 | 3.577 | 1.05 | 0.92 | 0.64 | | 0.67 | 0.60 | 0.78 | 1.70 | 2.17 | 33.69 |
| APC | 0.00 | 0.00 | 2.09 | 0.45 | 0.00 | 1.02 | 0.73 | 0.42 | 0.29 | 0.36 | 0.00 | 0.15 | 0.37 | 1.05 | 2.60 | 1.41 | 1.01 | 1.35 | 0.79 | 0.93 | 0.00 | 1.95 | 1.30 | 1.06 | 0.00 | 0.00 | 0.79 | 2.30 | 0.51 | | 2.72 | 1.91 | 1.44 | 0.00 | 29.01 |
| SNIR 685 | 0.00 | 3.17 | 0.00 | 0.00 | 1.04 | 0.00 | 0.00 | 0.00 | 0.00 | 1.78 | 0.00 | 8.14 | 1.31 | 4.376 | 2.30 | 0.00 | 0.00 | 4.086 | 1.38 | 6.357 | 0.00 | 0.00 | 3.409 | 0.00 | 4.69 | 0.00 | 0.00 | 0.00 | 2.46 | 1.50 | | 3.526 | 2.72 | 2.22 | 54.47 |
| APCR700 | 1.54 | 0.94 | 0.00 | 0.00 | 0.99 | 1.39 | 2.44 | 1.31 | 0.67 | 0.84 | 0.54 | 3.516 | 1.63 | 1.27 | 1.40 | 2.28 | 2.20 | 2.44 | 2.47 | 1.73 | 1.22 | 1.59 | 2.53 | 2.17 | 1.93 | 1.41 | 1.03 | 0.87 | 0.84 | 3.503 | 3.623 | | 4.138 | 2.01 | 56.47 |
| APC750 | 2.13 | 1.18 | 3.71 | 1.66 | 1.31 | 0.03 | 0.96 | 0.00 | 1.50 | 1.59 | 1.45 | 5.576 | 0.00 | 2.88 | 0.00 | 2.16 | 0.86 | 0.93 | 3.85 | 0.00 | 0.00 | 1.11 | 0.00 | 1.98 | 3.739 | 0.00 | 1.55 | 0.74 | 1.29 | 2.93 | 2.24 | 0.76 | | 4.596 | 52.71 |
| APC810 | 0.00 | 0.85 | 0.00 | 0.00 | 0.00 | 0.01 | 0.00 | 1.52 | 0.52 | 0.03 | 0.00 | 0.00 | 0.76 | 0.00 | 0.00 | 0.00 | 1.04 | 2.24 | 0.00 | 1.07 | 0.02 | 0.00 | 0.49 | 0.56 | 0.00 | 0.00 | 0.73 | 0.01 | 0.76 | 1.37 | 0.68 | 0.89 | 2.64 | | 16.21 |
| Sum | 24.08 | 44.04 | 122.66 | 54.14 | 19.67 | 24.76 | 28.59 | 28.36 | 19.50 | 38.01 | 31.07 | 146.75 | 41.26 | 38.65 | 30.04 | 36.39 | 34.42 | 41.72 | 54.88 | 61.73 | 37.25 | 34.21 | 44.07 | 30.00 | 63.22 | 39.02 | 37.39 | 29.06 | 19.93 | 35.51 | 35.09 | 30.63 | 35.82 | 28.95 | 1420.87 |

*Supplementary Figure 7 Spillover spreading matrix of healthy donor cells stained with the panel. Some of the spread into the detectors for BUV496 and BV510 likely originates from residual autofluorescence rather than fluorochrome-emitted signal, since many of the fluorochromes that appear to emit it have longer peak emission wavelengths than these detectors and do not emit in the 520nm - 550nm UV and violet regions.*

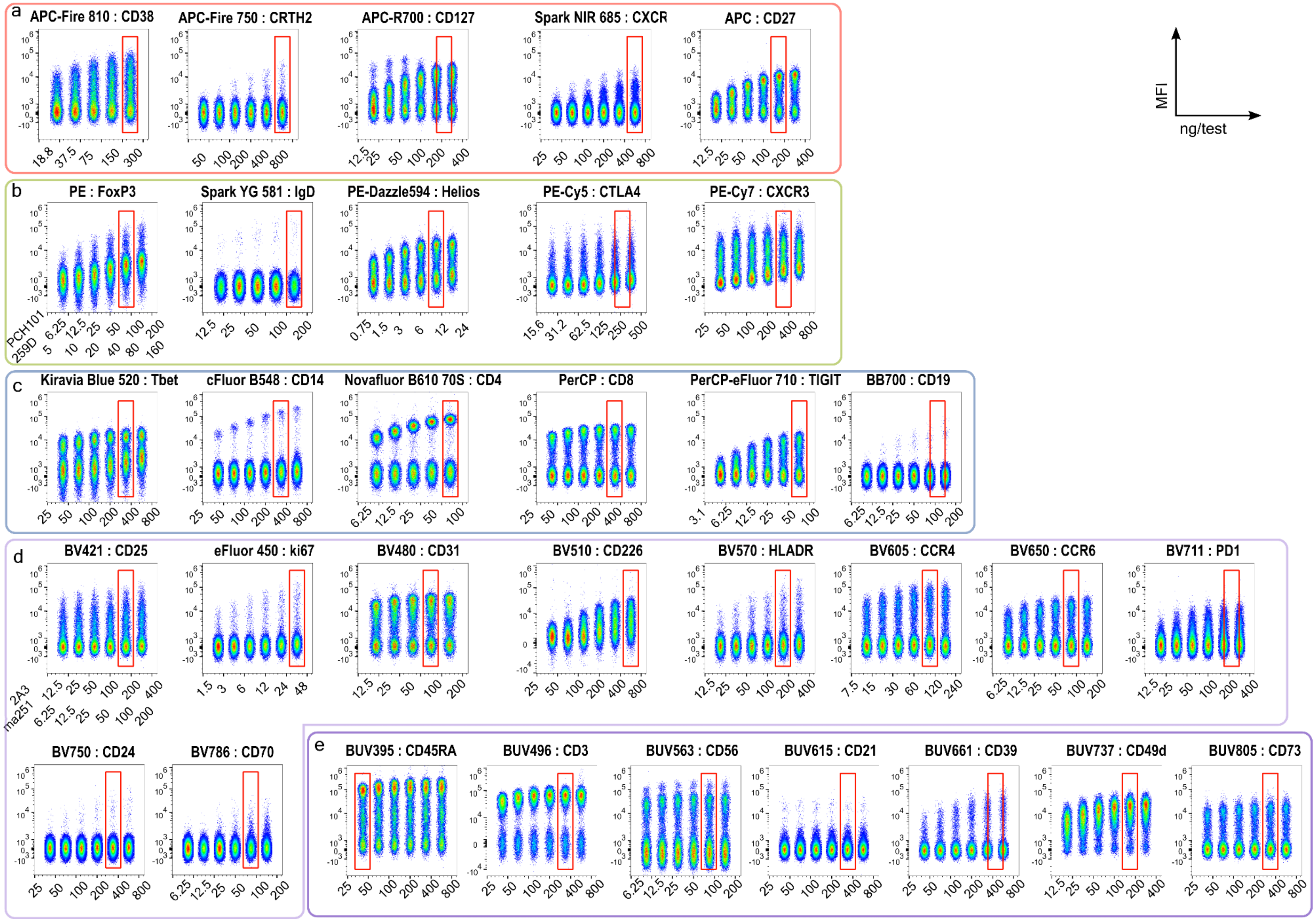


*Supplementary Figure 8 Titrations for antibody-fluorochrome conjugates primarily excited by the red (a), yellow-green (b), blue (c), violet (d) and UV (e ) lasers, determined by serial dilutions ranging from 1:25 or 1:50 to 1:1600 on 200,000 cells/sample. Equal event numbers were acquired for each sample. Live single cells were exported and concatenated in FlowJo v10.8.1. Optimum concentration is indicated by a red rectangle (in some cases, this is the median concentration between two dilutions). Optimum concentration was selected to maximise staining index, and in some cases- increased 1.5-fold (to ensure saturation when many millions of cells are stained, e.g. CD39 BUV661) or decreased (to limit data spread or background staining, e.g. CD19 BB700). Concentration is expressed in ng/100µL test. See Supplementary Table 4.*

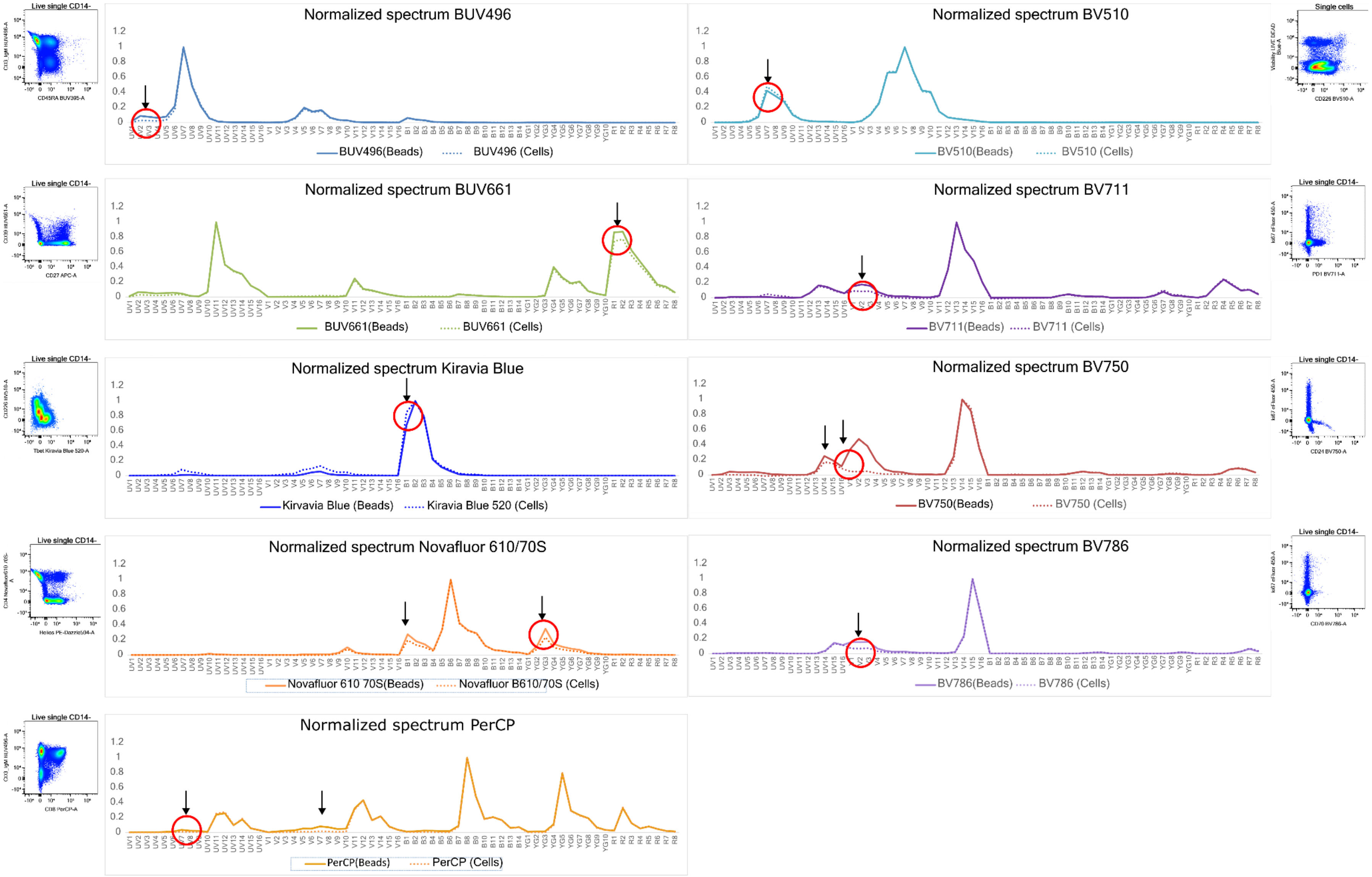


*Supplementary Figure 9 Comparison of unmixing results using beads or cells as reference controls. Unmixing was performed in SpectroFlo v3.0 and single-stain control spectra were normalised to the maximum peak emission. Fluorochromes which can only be unmixed using cells are shown. Arrows indicate discrepancies in the normalised spectra when unmixing with beads is performed. Inset scatter plots illustrate potential unmixing errors due to these discrepancies. All other fluorochromes can be unmixed successfully using UltraComp beads.*

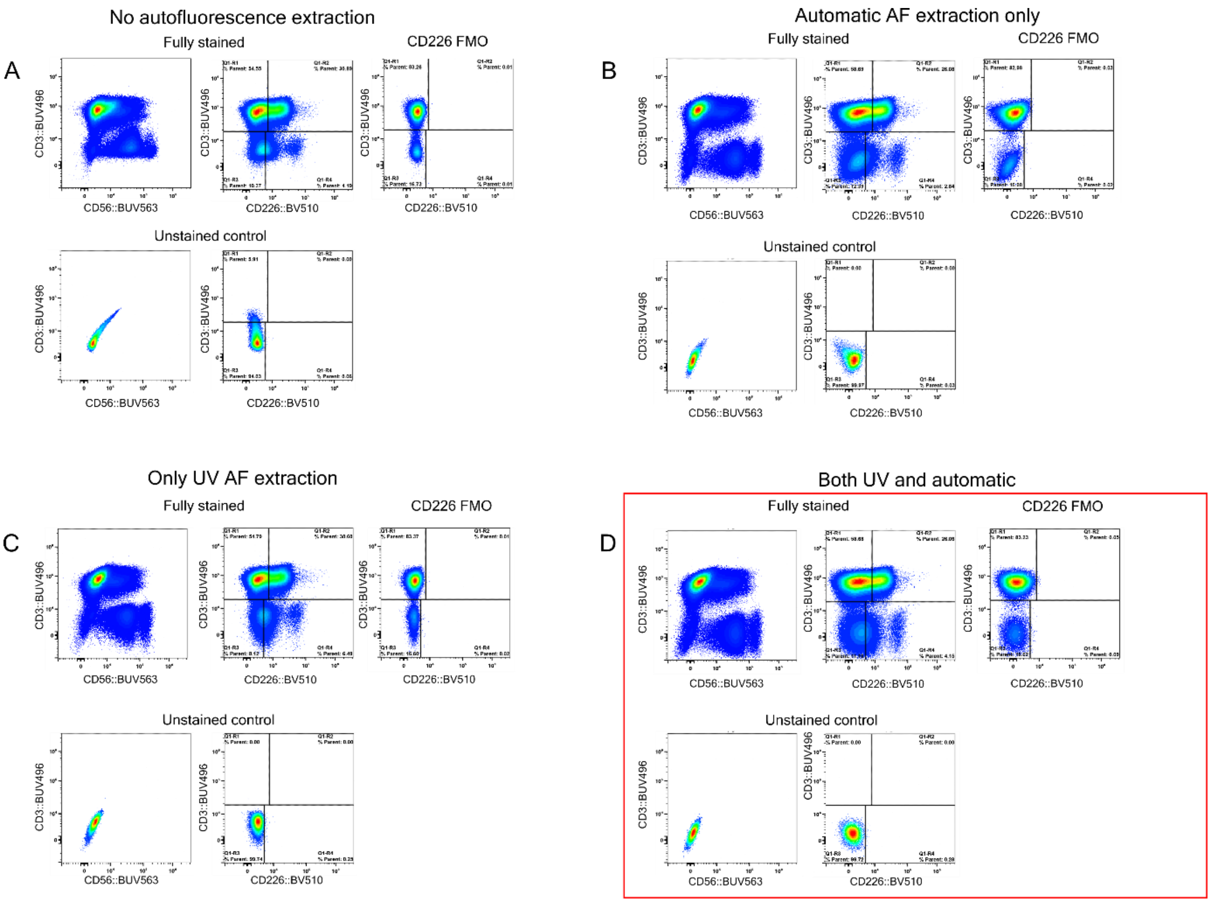


*Supplementary Figure 10 Comparison of unmixing methods to remove autofluorescence, which is evident as positive signal in BUV496 and BUV563 on unstained samples and in the negative population of multi-stained samples. A- Unmixing without any AF extraction has the highest AF signal. B- Unmixing using the Automatic AF extraction in SpectroFlow v3.0 does not resolve this issue. C- Unmixing where AF is assigned a "virtual" fluorochrome as an additional reference control. Single lymphocytes were gated in raw FCS files of unstained controls acquired with each experiment. The events with the highest signal in channel UV7 were exported as a new FCS file (called "UV AF") and used as a new reference control to perform unmixing. The negative population for this control is total unstained lymphocytes. D- The optimum selected method in SpectroFlo v3.0 was a combination of B and C (to remove AF from the UV and violet spectrum). Other software, or updated versions of SpectroFlo, may use different unmixing algorithms, and therefore would require testing of optimal unmixing.*

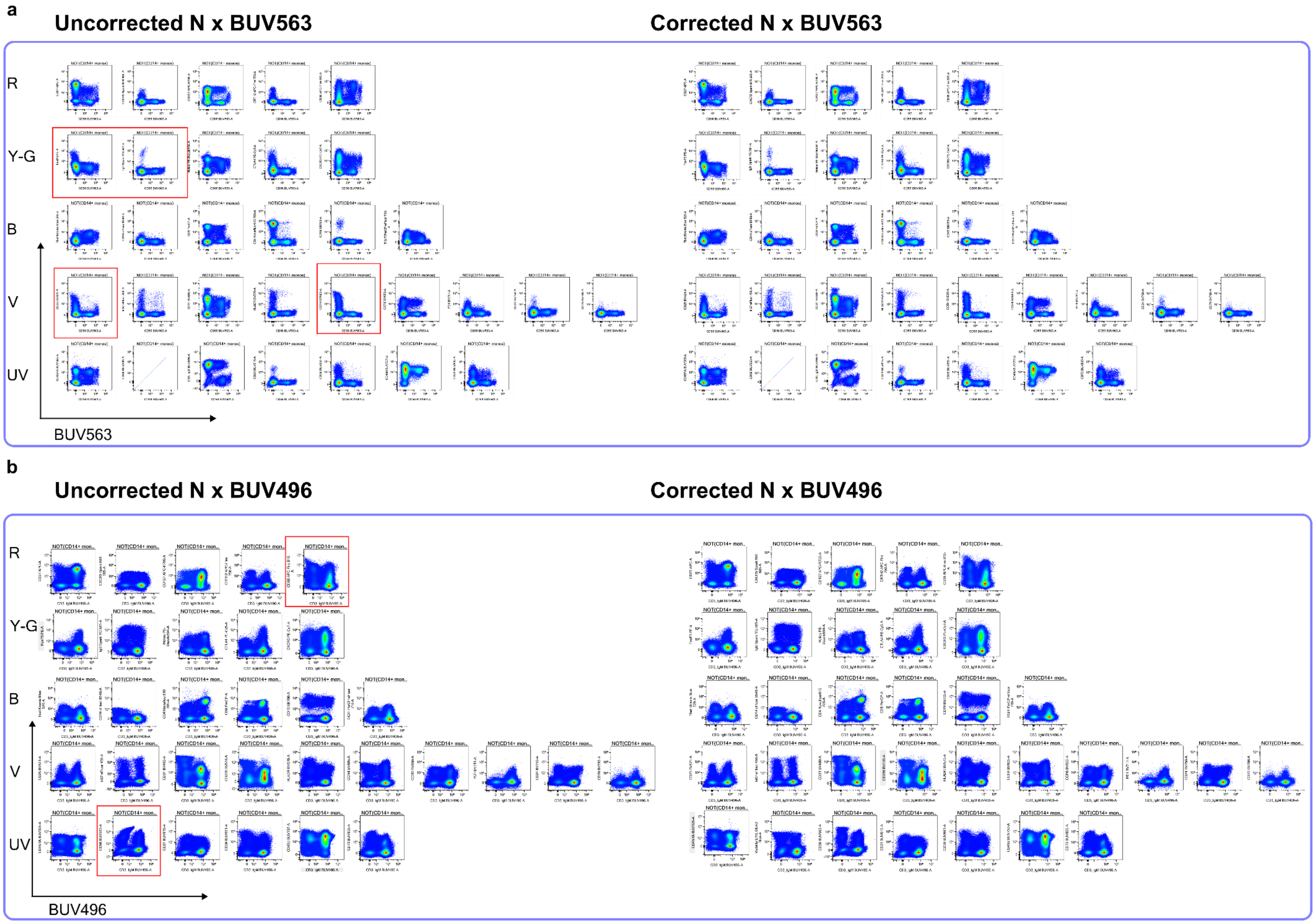


*Supplementary Figure 11 Unmixing accuracy of the multi-stained sample. Example spillover corrections for two fluorochromes, BUV563 (a) and BUV496 (b). N x fluorochrome plots display CD14- single cells, cleaned of debri and aggregates. Plots are arranged according to primary laser excitation, from long to short wavelength. Plots requiring correction are highlighted in red boxes*

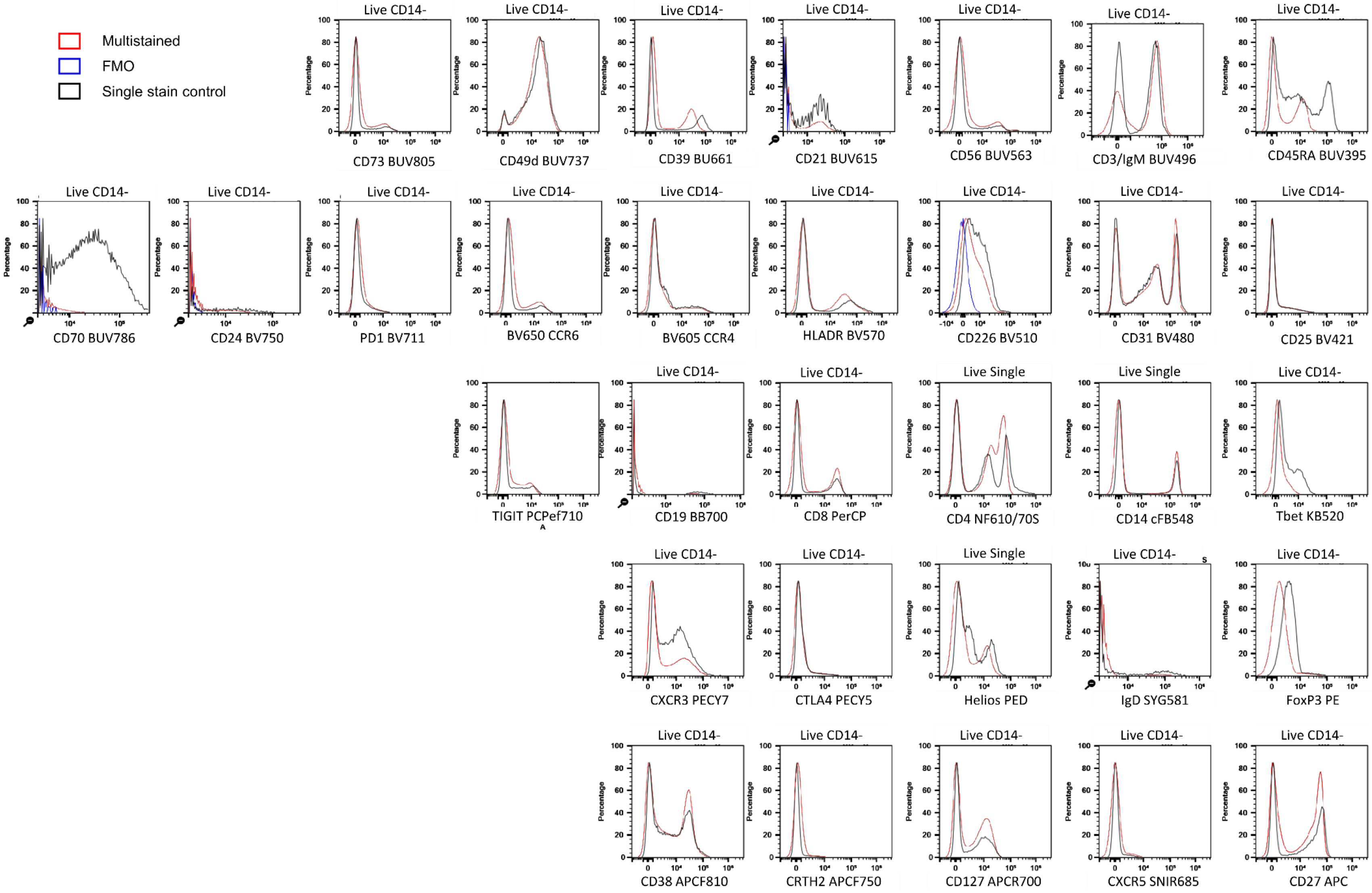


*Supplementary Figure 12 Multi-stained (red) and single-stained (black) samples from the same donor were compared to evaluate resolution for each marker. Live single CD14- cells are shown. For CD226 and CD70, FMO control samples (blue) are also shown to demonstrate positive expression more clearly. The negative population has a similar distribution in SS and MS samples, showing that spread has minimal impact on resolution. TBET signal appears less well resolved in this donor, who had unusually low expression confined to NK cells and some CD8 T-cells. Other donors expressing higher TBET are shown in Supplementary Figure 15 A.*

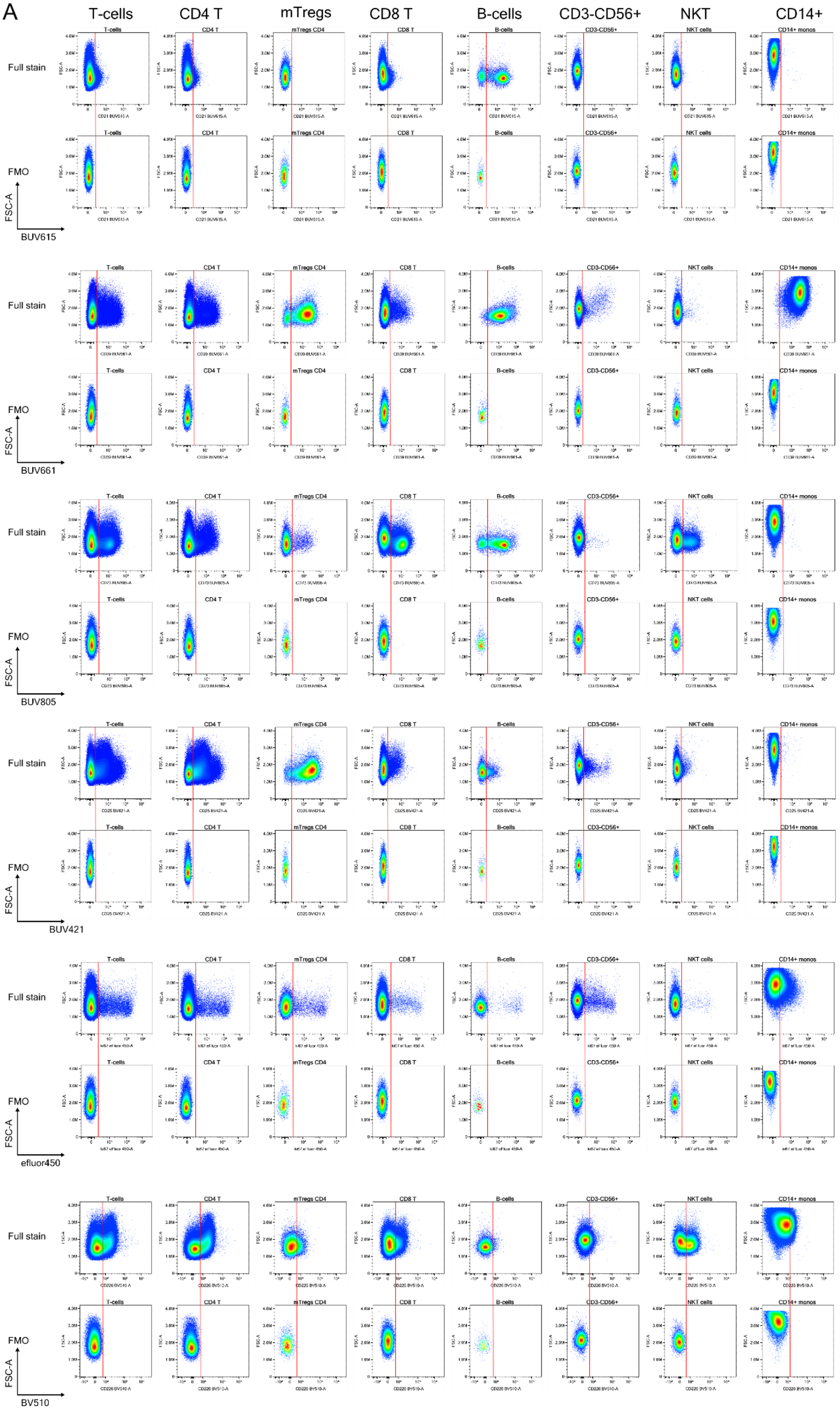
A
T-cells
CD4 T
mTregs
CD8 T
B-cells
CD3-CD56+
NKT
CD14+
Full stain
FMO
FSC-A
BUV615
BUV661
BUV805
BUV421
efluor450
BV510

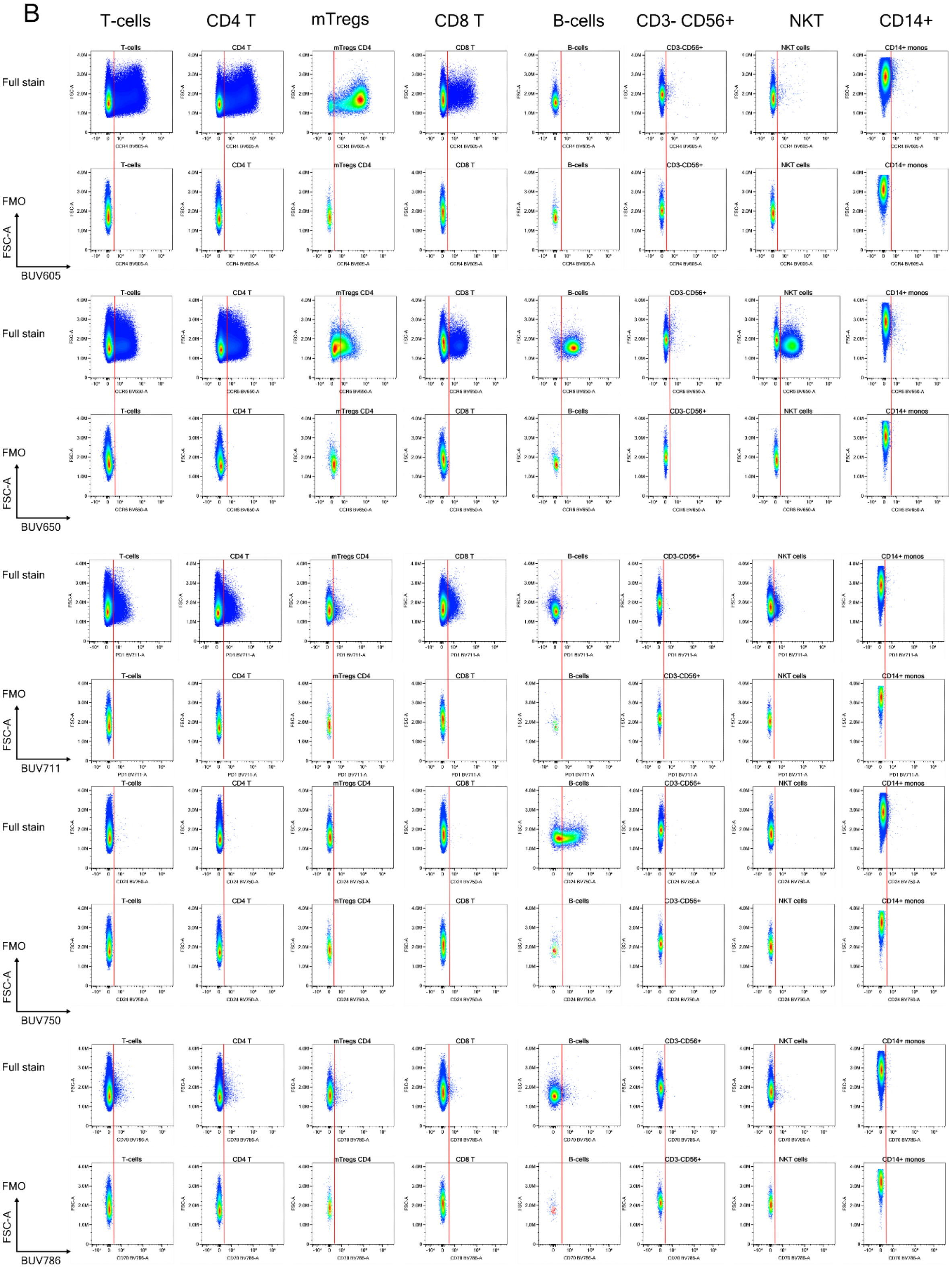
B
T-cells
CD4 T
mTregs
CD8 T
B-cells
CD3- CD56+
NKT
CD14+
Full stain
FMO
FSC-A
BUV605
Full stain
FMO
FSC-A
BUV650
Full stain
FMO
FSC-A
BUV711
Full stain
FMO
FSC-A
BUV750
Full stain
FMO
FSC-A
BUV786

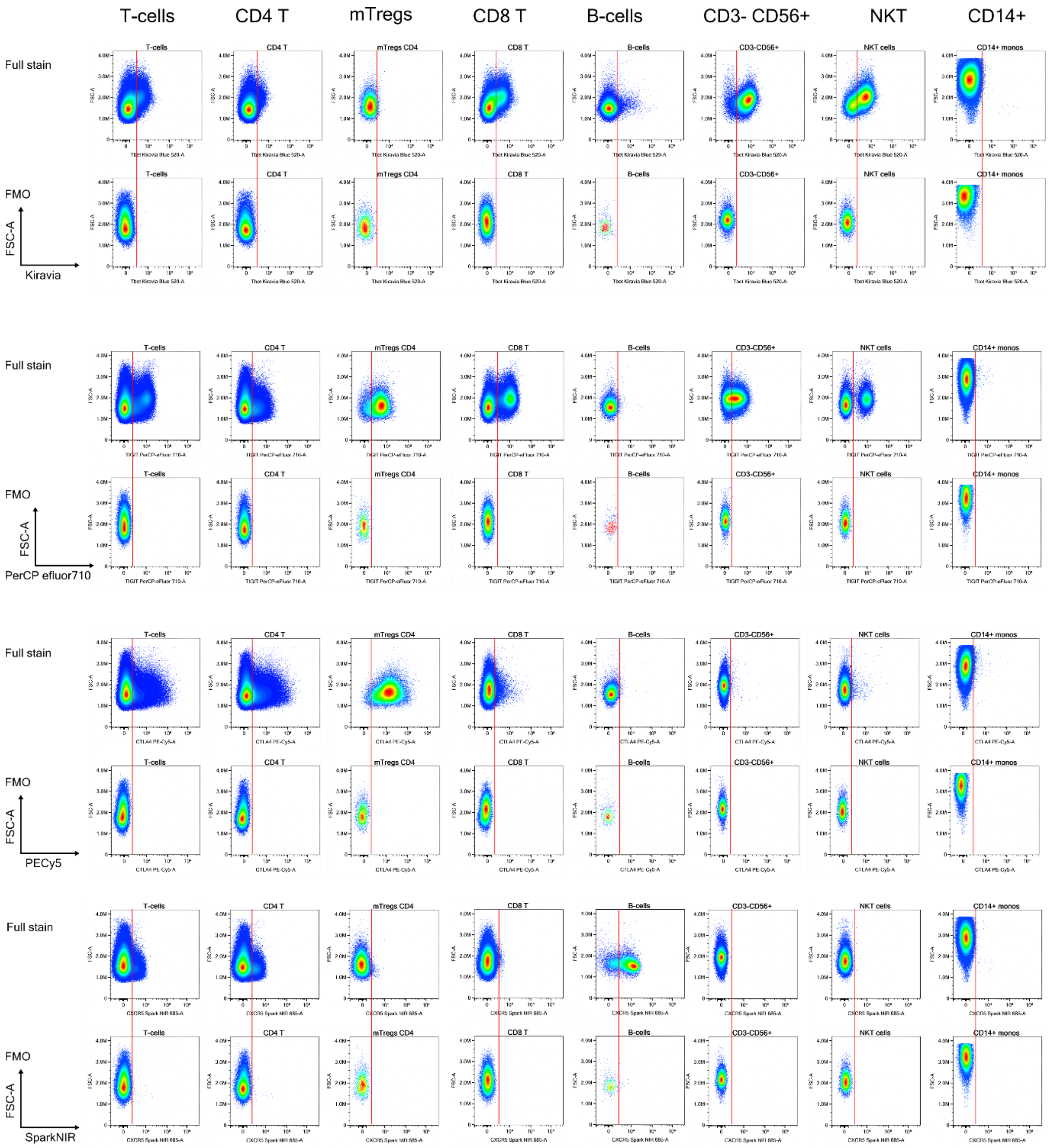


*Supplementary Figure 13 FMO controls for the markers shown (A-C) were used to set positive and negative gates and confirm good resolution of these markers on each of the populations shown (bulk T-cells, CD4, CD8 T-cells, CD4 Tregs, B-cells, NK cells and NKT cells. CD14+ monocytes are shown but they do not express the majority of the markers on this panel, which was targeted at lymphocytes.*

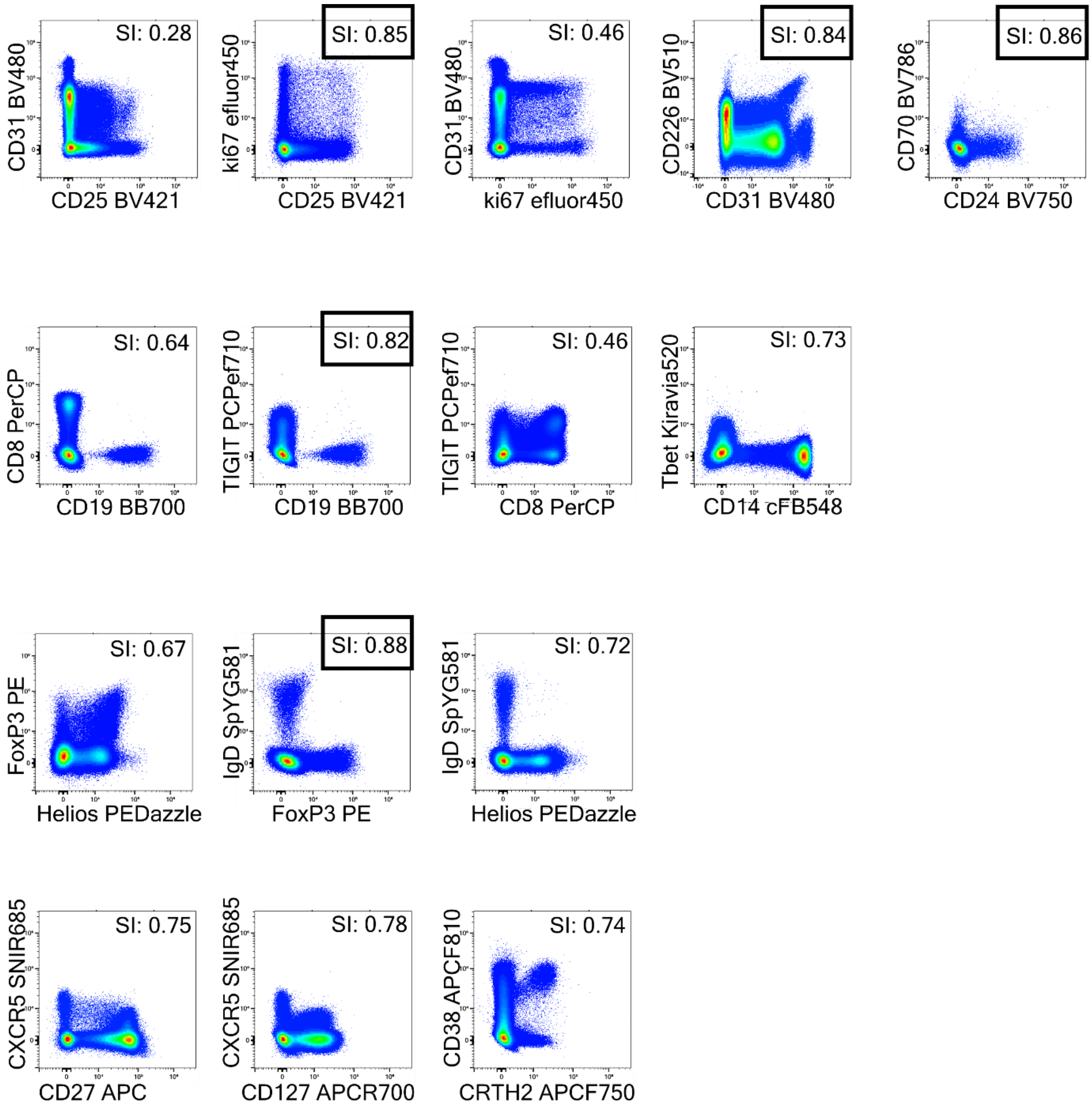


*Supplementary Figure 14 Impact of highly similar fluorochromes on resolution was evaluated in NxN plots of live single CD14- cells. The most similar pairs are shown here, and each marker remains well resolved.*

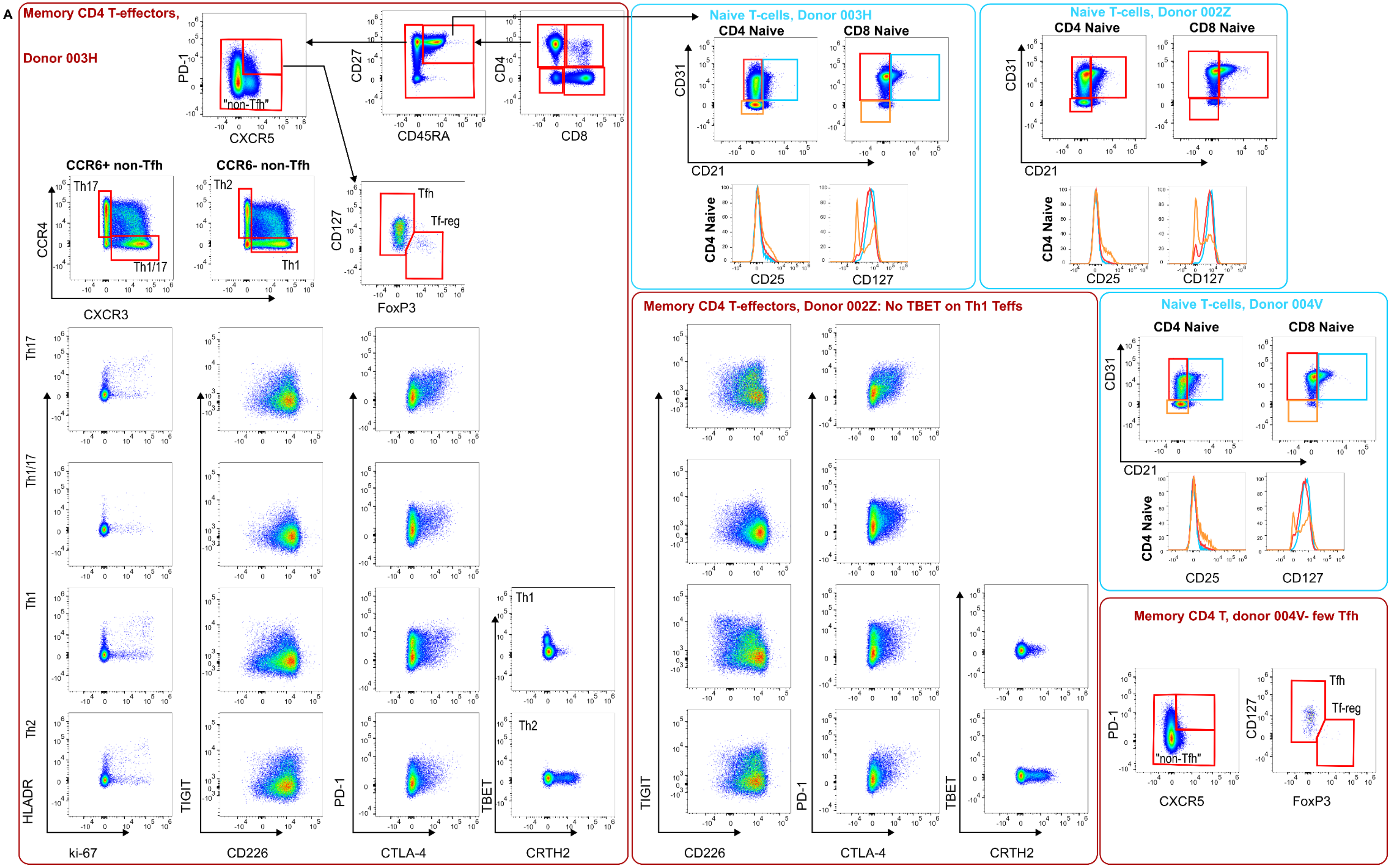
A
Memory CD4 T-effectors, Donor 003H
PD-1
CXCR5
"non-Tfh"
CD27
CD45RA
CD4
CD8
CCR6+ non-Tfh
CCR6- non-Tfh
Th17
Th1/17
Th2
Th1
CCR4
CXCR3
CD127
FoxP3
Tfh
Tf-reg
HLADR
ki-67
TIGIT
CD226
PD-1
CTLA-4
TBET
CRTH2
Naive T-cells, Donor 003H
CD4 Naive
CD8 Naive
CD31
CD21
CD25
CD127
Naive T-cells, Donor 002Z
CD4 Naive
CD8 Naive
CD31
CD21
CD25
CD127
Memory CD4 T-effectors, Donor 002Z: No TBET on Th1 Teffs
TIGIT
CD226
PD-1
CTLA-4
TBET
CRTH2
Naive T-cells, Donor 004V
CD4 Naive
CD8 Naive
CD31
CD21
CD25
CD127
Memory CD4 T, donor 004V- few Tfh
PD-1
CXCR5
"non-Tfh"
CD127
FoxP3
Tfh
Tf-reg

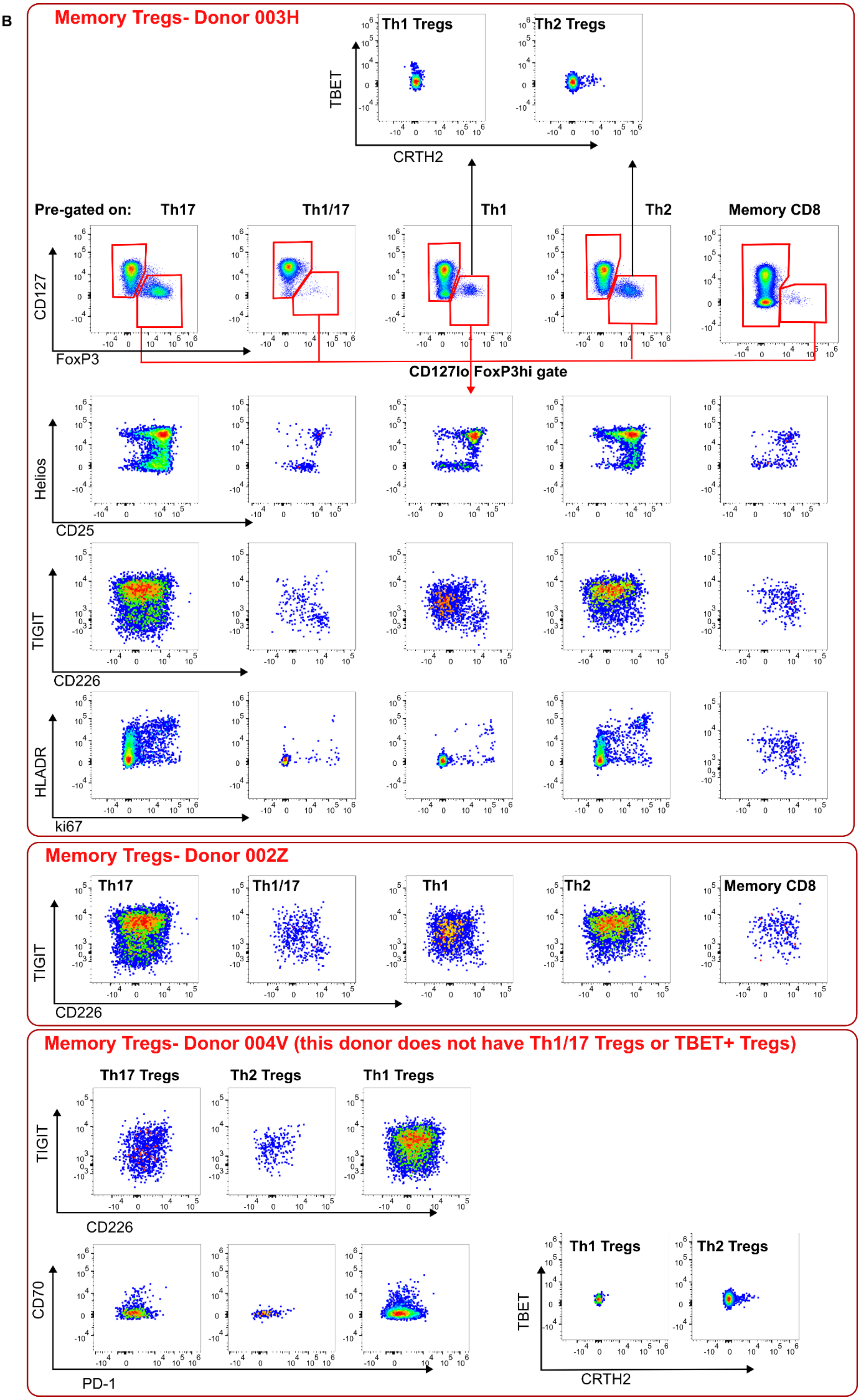
B
Memory Tregs- Donor 003H
Th1 Tregs
Th2 Tregs
TBET
CRTH2
Pre-gated on:
Th17
Th1/17
Th1
Th2
Memory CD8
CD127
FoxP3
CD127lo FoxP3hi gate
Helios
CD25
TIGIT
CD226
HLADR
ki67
Memory Tregs- Donor 002Z
Th17
Th1/17
Th1
Th2
Memory CD8
TIGIT
CD226
Memory Tregs- Donor 004V (this donor does not have Th1/17 Tregs or TBET+ Tregs)
Th17 Tregs
Th2 Tregs
Th1 Tregs
TIGIT
CD226
CD70
PD-1
Th1 Tregs
Th2 Tregs
TBET
CRTH2

C

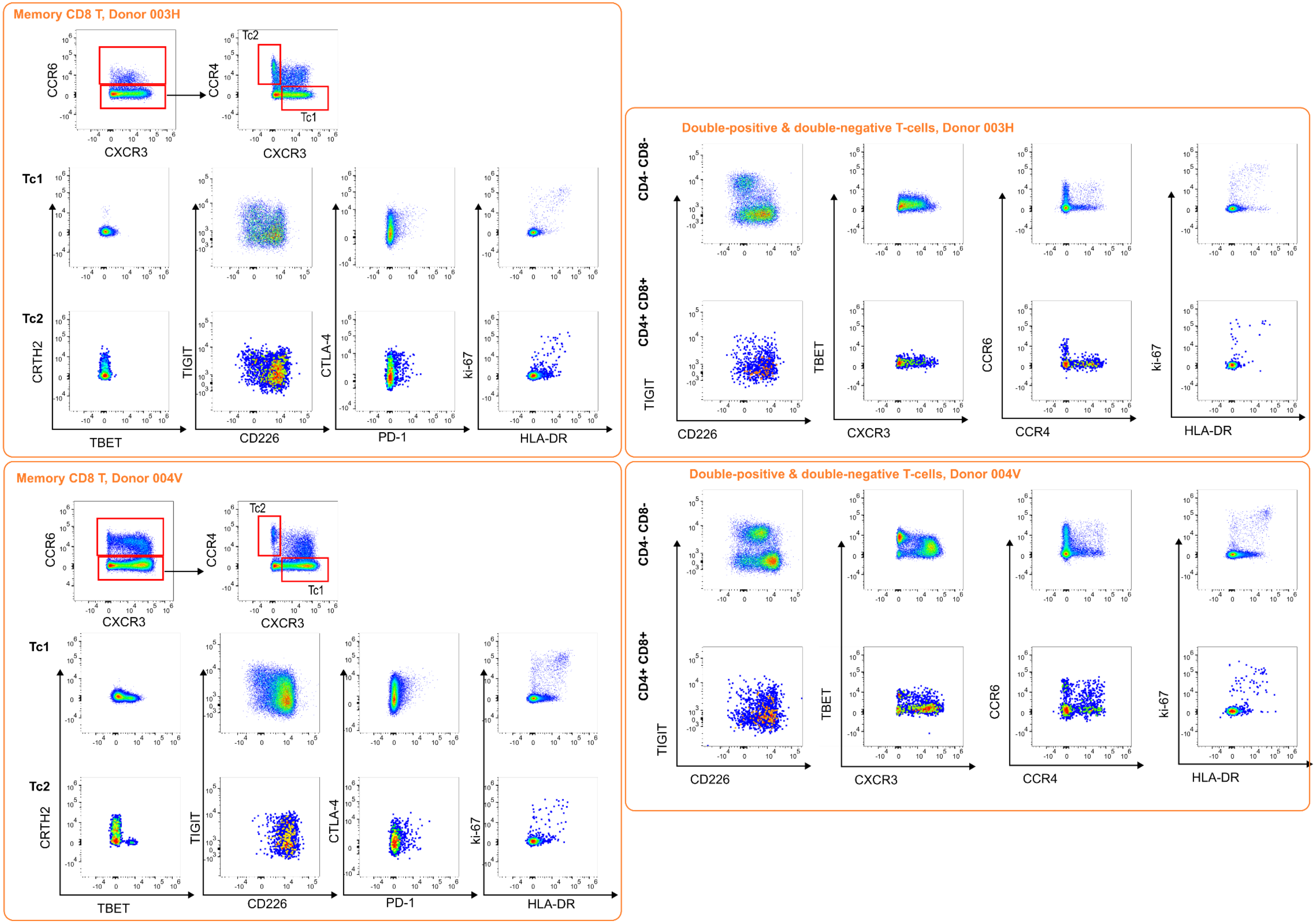
Memory CD8 T, Donor 003H
CCR6
CXCR3
CCR4
Tc2
Tc1
Tc1
Tc2
CRTH2
TBET
TIGIT
CD226
CTLA-4
PD-1
ki-67
HLA-DR
Double-positive & double-negative T-cells, Donor 003H
CD4- CD8-
CD4+ CD8+
TIGIT
CD226
TBET
CXCR3
CCR6
CCR4
ki-67
HLA-DR
Memory CD8 T, Donor 004V
CCR6
CXCR3
CCR4
Tc2
Tc1
Tc1
Tc2
CRTH2
TBET
TIGIT
CD226
CTLA-4
PD-1
ki-67
HLA-DR
Double-positive & double-negative T-cells, Donor 004V
CD4- CD8-
CD4+ CD8+
TIGIT
CD226
TBET
CXCR3
CCR6
CCR4
ki-67
HLA-DR

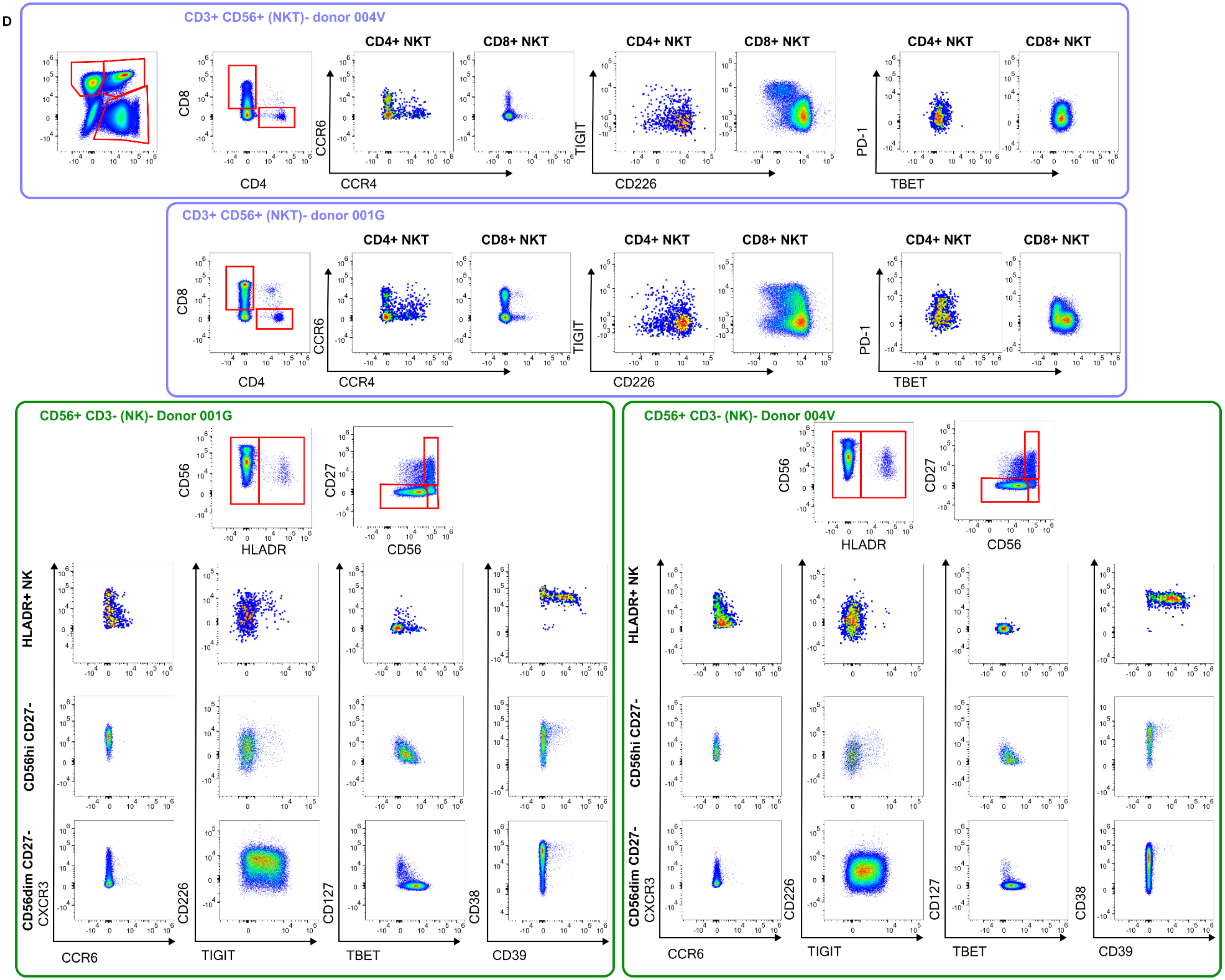
D
CD3+ CD56+ (NKT)- donor 004V
CD4+ NKT
CD8+ NKT
CD4+ NKT
CD8+ NKT
CD4+ NKT
CD8+ NKT
CD8
CD4
CCR6
CCR4
TIGIT
CD226
PD-1
TBET
CD3+ CD56+ (NKT)- donor 001G
CD4+ NKT
CD8+ NKT
CD4+ NKT
CD8+ NKT
CD4+ NKT
CD8+ NKT
CD8
CD4
CCR6
CCR4
TIGIT
CD226
PD-1
TBET
CD56+ CD3- (NK)- Donor 001G
CD56
HLADR
CD27
CD56
HLADR+ NK
CD56hi CD27-
CD56dim CD27-
CXCR3
CCR6
CD226
TIGIT
CD127
TBET
CD38
CD39
CD56+ CD3- (NK)- Donor 004V
CD56
HLADR
CD27
CD56
HLADR+ NK
CD56hi CD27-
CD56dim CD27-
CXCR3
CCR6
CD226
TIGIT
CD127
TBET
CD38
CD39

E

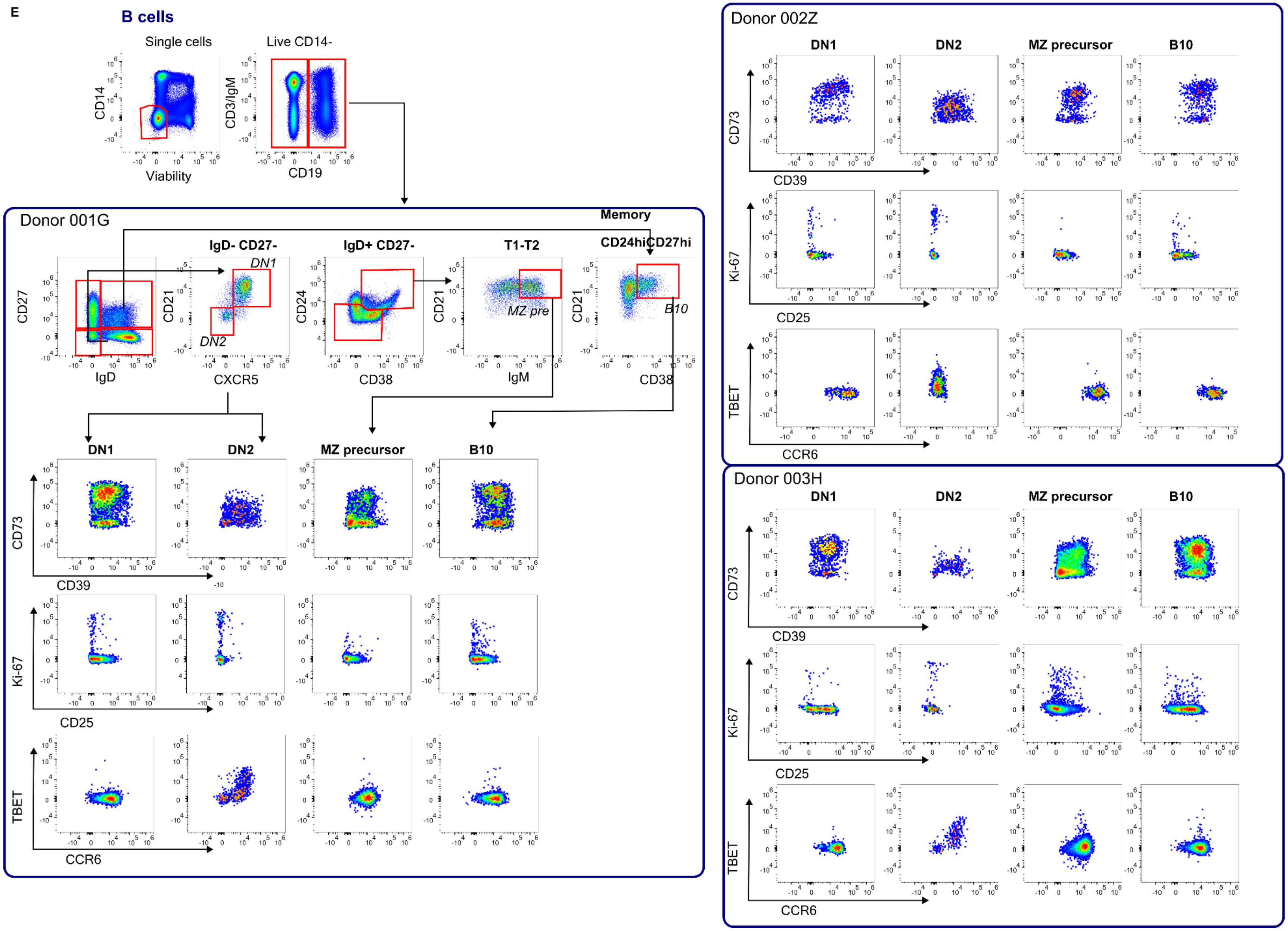


*Supplementary Figure 15: Examples of differences between donors stained with this panel on: CD4 T-effector (A), CD4 Treg (B), CD8 T-effector and double-positive/double-negative T-cells (C), NK/NKT cells (D) and B-cells (E ).*

| Supplementary Table 1: Expression and roles of markers in T-cells | | |
|---|---|---|
| **Specificity** | **Expression and role in peripheral blood** | **Example functional relevance** |
| CD21 | **Treg -** **CD4 Teff +** **CD8 Teff +**<br>Expressed on the most recent RTE CD4 and CD8 Teffs, but not Tregs (37)<br>Declines with homeostatic proliferation(38). | • Imbalanced Treg : Teff thymopoiesis is associated with development of GvHD after HSCT (39) |
| CD31 | **Treg +++** **CD4 Teff +++** **CD8 Teff +++**<br>Expressed on naïve RTE CD4 Teff and Tregs. Declines with homeostatic proliferation.<br>Does not indicate CD8 RTEs, unlike CD21. | • RTE Tregs are specifically reduced in MS.(40)<br>NB: IL-7 maintains CD31 expression, so CD31 is not a suitable RTE marker in a high-IL-7 context. |
| CD39 | **Treg +++** **CD4 Teff +** **CD8 Teff +**<br>Catalyses conversion of extracellular ATP to AMP, a first step in the production of immunosuppressive adenosine.<br>Highly expressed on CCR6+ Tregs (41). | • CD39+ Tregs are deficient in RRMS(42).<br>• CD39+ Tregs make adenosine upon contact with cancer-derived CD73+ exosomes.(43)<br>• Circulating exhausted CD39+ PD1+ CD4 Teffs predict response to ICB in cancer(44). |
| CD73 | **Treg ++** **CD4 Teff +** **CD8 Teff +**<br>Only co-expressed with CD39 on human Tregs following activation.<br>Also expressed on some naïve and memory Teffs. | • Human Tregs gain CD73 after repeated polyclonal *in vitro* expansion, becoming super-suppressive(35).<br>• Fewer circulating CD8+ CD73+ PD1+ T-cells correlate with better response to nivolumab in melanoma(45). |
| CD49d | **Treg +** **CD4 Teff +++** **CD8 Teff +++**<br>Integrin α4 subunit, allows access to brain (α4β1) and gut (α4β7). | • (Murine) Teffs, but not Tregs, use CD49d to enter the CNS.(46)<br>• CD49d is the target of natalizumab, a drug used in MS. |
| CD226 | **Treg +** **CD4 Teff +++** **CD8 Teff +++**<br>Competes with TIGIT for the same ligand, CD112/CD155.<br>Predominantly expressed on Teffs, but also found on Tregs. | • CD226+ CD127-CD25+ T-cells can produce inflammatory cytokines. CD226 exclusion produces purer more stable Tregs for therapeutic expansion.(48)<br>• |
| CD226 | CD226 gene is a risk locus in multiple autoimmune diseases(47). | • Treg specific deletion of CD226 in NOD mice maintains Treg stability and reduces diabetes.(49)<br>Human CD226- CD8 Teffs are hyporesponsive. Loss of CD226 in mouse tumour models limits response to ICB.(50) |

**(continued) Supplementary Table 1: Expression and roles of markers in T-cells**

| | | |
|---|---|---|
| TIGIT | **Treg +++ CD4 Teff + CD8 Teff**<br>TIGIT competes with CD226 for CD112/155, and provides a co-inhibitory, suppressive signal.<br>Expression in Tregs correlates with phenotypic stability. In Teffs, TIGIT signifies exhaustion. | • High TIGIT:CD226 ratio in Tregs favours Treg infiltration in melanoma(51).<br>• TIGIT stimulation corrects the functional defect of Th1-like Tregs in MS(52). |
| PD-1 | **Treg +++ CD4 Teff + CD8 Teff+**<br>PD-1 provides an inhibitory signal.<br>Expression on Teff populations indicates exhaustion. | • Circulating PD-1+ Tregs are dysfunctional in glioblastoma(53).<br>• PD-1 is required for Treg survival during IL-2 induced therapeutic expansion *in vivo*.(54) |
| CD70 | **Treg + CD4 Teff + CD8 Teff+**<br>CD70 is expressed on a very small proportion of Tregs, reciprocally to CD27, its ligand. | • Tregs gain CD70 expression after repeated polyclonal expansion *in vitro*, becoming less suppressive because they can provide co-stimulation to Teffs.(55) |
| CTLA-4 | **Treg +++ CD4 Teff + CD8 Teff +**<br>A major cell-contact dependent suppressive mechanism.<br>Competes with CD28 and removes CD80/CD86 from APCs by transendocytosis. | • IL-2 therapy increases CTLA-4 expression by Tregs (56)<br>• CTLA-4 may control human Teff motility and contact time with APCs (57)<br>• CTLA-4 inhibition is an established ICB cancer therapy. |

*Supplementary Table 1 Roles of selected markers on the panel in T-effectors and Tregs, and non-exhaustive examples of involvement in human autoimmunity or cancer.*

**Supplementary Table 2: Expression and roles of markers in B-cells**

| Specificity | Role and expression in peripheral blood | Example functional relevance |
|---|---|---|
| CD39 and CD73 | CD39+ B cells suppress CD4 Teff responses via adenosine production (58). Up to 90% of human B-cells co-express CD39 and CD73. | Adenosine production by CD73+ B-cells is reduced in SLE.(59) |
| CXCR3, CXCR5 and CCR6 | Regulation of cell trafficking. | CXCR3+ autoreactive B-cells are present in individuals with RA(60). Expression and roles in autoimmune disease are relatively underexplored. |
| TBET | Regulation of immunoglobulin class-switching | TBET+ CD27-IgD- B-cells, also known as DN2, develop directly from naïve B-cells, produce autoantibodies and are enriched in SLE.(61) |

*Supplementary Table 2 Roles of selected markers on the panel in the B-cell compartment. RA= rheumatoid arthritis, MS= multiple sclerosis, SLE= systemic lupus erythematosus. Non-exhaustive examples for the involvement of these markers in autoimmunity or inflammation are shown, preferably in the peripheral blood compartment. CD49d, CD226, CD31, CD70, are also expressed but understanding of their roles in B-cells is still relatively limited, and therefore not listed.*

**Supplementary Table 3: Expression and roles of markers in NK cells**

| Specificity | Role and expression in healthy control blood | Example functional relevance |
|---|---|---|
| CD38 | Expressed equally by CD56bright and CD56dim NK cells. | Adenosine production by healthy CD56$^{bright}$ NK cells limits CD4 T-cell proliferation (62). |
| TIGIT | Inhibitory receptor and exhaustion marker. TIGIT expression in circulating NK cells correlates with lower cytotoxic potential in healthy individuals(63). Intrahepatic CD56$^{bright}$ NK cells express more TIGIT than their circulating counterparts(64), i.e. expression may be relatively limited in blood. | TIGIT blockade reverses (mouse) NK exhaustion to promote tumour cytotoxicity (65) |
| CD226 | Activating receptor, promotes cytotoxic function. It is expressed on both CD56bright and CD56dim NK cells. | Human CD226- NK cells have limited killing ability and produce little IFNγ.(66) |
| PD-1 | Inhibitory receptor. Expressed on NK cells in up to 25% of healthy human subjects. Suppresses NK function (67). | PD-1+ NK cells have reduced cytotoxic and cytokine-producing potential in PD1L+ tumour environment (68). |
| CXCR3 | Predominantly expressed on CD56$^{bright}$ NK cells, and likely controls trafficking to tissue (e.g. skin and liver) | CXCR3+ CD56$^{bright}$ NK cells have anti-fibrotic properties in chronic liver disease (69). CXCR3+ CD56$^{bright}$ NK cells are proinflammatory in psoriasis.(70) |
| TBET | Regulates the expression of genes involved in cytotoxicity and IFNγ production. | CRISPR/Cas9 edited TBET- human NK cells are phenotypically unstable and fail to control tumour growth (71) |

*Supplementary Table 3 Roles of selected markers on the panel in NK cell function, and their example relevance to human disease. Other markers, e.g. CD49d, CD39, CD73 are also expressed, but not listed due to relatively limited understanding of their roles.*

| Antibodies and optimum amounts used in this panel | | | | | | |
|---|---|---|---|---|---|---|
| **Antibody** | **Clone** | **Catalogue #, Isotype** | **Supplier** | **Stock (ng/ml)** | **Volume (µL/test)** | **Amount (ng/test)** |
| **Surface staining part 1 (21°C)** | | | | | | |
| CCR4 BV605 | L291H4 | 359418, IgG1, κ | Biolegend | 60000 | 2 | 120 |
| CCR6 BV650 | G034E3 | 353426, IgG2b,k | Biolegend | 50000 | 2 | 100 |
| CXCR3 PECy7 | G025H7 | 353720, IgG1,k | Biolegend | 200000 | 2 | 400 |
| CXCR5 SparkNIR685 | J252D4 | 356947, IgG1,k | Biolegend | 200000 | 4 | 800 |
| **Surface staining part 2 (4°C)** | | | | | | |
| CD45RA BUV395 | HI100 | 740296, IgG1, κ | BD | 200000 | 0.125 | 25 |
| CD3 BUV496 | SK7 | 741206, IgG1, κ | BD | 200000 | 2 | 400 |
| IgM BUV496 | UCHB1 | 750366, IgG1, κ | BD | 200000 | 4 | 800 |
| CD21 BUV615 | B-ly4 | 751413, IgG1, κ | BD | 200000 | 2 | 400 |
| CD56 BUV563 | NCAM16.2 | 612929, IgG2b, κ | BD | 50000 | 2 | 100 |
| CD39 BUV661 | TU66 | 749967, IgG2b, κ | BD | 200000 | 3 | 600 |
| CD49d BUV737 | G9F10 | 612850, IgG1, κ | BD | 100000 | 2 | 200 |
| CD73 BUV805 | AD2 | 748584, IgG1, κ | BD | 200000 | 2 | 400 |
| CD25 BV421 | MA251 | IgG1,k 356114 | Biolegend | 50000 | 2 | 100 |
| CD25 BV421 | 2A3 | 564033, IgG1, κ | BD | 100000 | 2 | 200 |
| CD31 BV480 | WM59 | 566195, IgG1, κ | BD | 100000 | 1 | 100 |
| CD226 BV510 | DX11 | 742494, IgG1, κ | BD | 200000 | 4 | 800 |
| CD226 BV510 | 11A8* | 338330, IgG1k | Biolegend | 200000 | 3 | 600 |
| HLADR BV570 | L243 | 307638, IgG2a,k | BD | 100000 | 2 | 200 |
| PD-1 BV711 | EH12.1 | 564017, IgG1, κ | BD | 100000 | 3 | 300 |
| CD24 BV750 | ML5 | 746890, IgG2a, κ | BD | 200000 | 2 | 400 |
| CD70 BV786 | Ki-24 | 565338, IgG3, κ | BD | 50000 | 2 | 100 |
| GITR BB515 | V27-580* | 567781, IgG2b, κ | BD | 200000 | 4 | 800 |
| CD14 cFluorB548 | 63D3 | R720115, IgG1,k | CytekBio | 200000 | 2 | 400 |
| CD4 NovaFluor610/70S | SK3 | H001T03B06, IgG1,k | ThermoFisher | 50000 | 2 | 100 |
| CD8 PerCP | SK1 | 344708, IgG1,k | Biolegend | 200000 | 2 | 400 |
| CD19 BB700 | SJ25C1 | 566396, IgG1, κ | BD | 50000 | 3 | 150 |
| TIGIT PerCPefluor710 | MBSA43 | 46950042, IgG1, k | eBioscience | 25000 | 4 | 100 |

| | | | | | | |
|---|---|---|---|---|---|---|
| IgD SparkYG581 | W18340F | IgG2,k 307805 | Biolegend | 100000 | 2 | 200 |
| CD27 APC | MT271 | 356410, IgG1,k | Biolegend | 100000 | 2 | 200 |
| CD127 APCR700 | HIL7RM21 | 565185, IgG1, κ | BD | 100000 | 3 | 300 |
| CRTH2 APCFire750 | BM16 | 350133, Rat IgG2a,k | Biolegend | 400000 | 2 | 800 |
| OX40 APCFire750 | BerACT35* | 350032, IgG1,k | Biolegend | 200000 | 4 | 800 |
| CD38 APCFire810 | HIT2 | 303549, IgG1,k | Biolegend | 150000 | 2 | 300 |
| **Intracellular** | | | | | | |
| Ki67 efluor450 | 20Raj1 | 48569942, IgG1, k | eBioscience | 12000 | 4 | 48 |
| TBET Kiravia Blue520 | 4B10 | 644838, IgG1,k | Biolegend | 200000 | 2 | 400 |
| FoxP3 PE | PCH101 | 12477642, IgG2a, k | eBioscience | 50000 | 2 | 100 |
| FoxP3 PE | 259D | 320208, IgG1,k | Biolegend | 40000 | 2 | 80 |
| HELIOS PE-Dazzle594 | 22F6 | 137232, Armenian Hamster IgG | BD | 6000 | 2 | 12 |
| CTLA-4 PECy5 | BNI3 | 555854, IgG2a, κ | BD | 25000 | 12.5 | 312.5 |
| **Selected antibodies used in panel development (not in final panel)** | | | | | | |
| Ki67 PECy5.5 | 20Raj1** | 355699-41, IgG1, k | eBioscience | 50000 | 0.1 | 5 |
| CD45RA PerCP | HI100** | 304156, IgG2b,k | Biolegend | 200000 | 5 | 1000 |

*Supplementary Table 4 Antibodies used in this panel. * Antibody used in customised version of the panel, see Supplementary Figure 2. ** Antibody used in panel development only.*
